# Dissociating volatility and stochasticity reveals transdiagnostic computational signatures of psychopathology

**DOI:** 10.64898/2026.05.22.727329

**Authors:** Xiaotong Fang, Payam Piray

## Abstract

Adaptive learning requires distinguishing volatility, changes in the latent state of the environment, from moment-to-moment stochasticity of observations. The two demand opposite adjustments to the learning rate: volatility calls for faster updating, stochasticity for slower. Disentangling them is computationally difficult because both inflate experienced variance, leaving the inference prone to systematic individual differences with potential consequences for psychopathology. Three computational phenotypes capture this variation: intact learners; stochasticity-blind learners, who over-update by treating noise as change; and volatility-blind learners, who under-update by treating change as noise. In two large online samples and across three tasks, we found a double dissociation between these phenotypes and transdiagnostic psychiatric dimensions: stochasticity-blind learners scored higher on Internalizing (anxiety, depression), volatility-blind learners on Externalizing (behavioral addiction, compulsivity).

Distinct symptom dimensions thus correspond to distinct failures of inference about uncertainty, supporting a selective rather than generalized account of learning-under-uncertainty deficits in psychopathology.

## Introduction

Behavior under uncertainty requires updating beliefs from incomplete, ambiguous, or shifting evidence, and this capacity is disrupted across many forms of psychopathology, including anxiety and depression, compulsivity-related disorders, behavioral addictions, and psychosis-related conditions ^1–6^. Anxiety disorders, for instance, often involve pronounced intolerance of uncertain situations, leading to excessive worry and avoidance, whereas behavioral addictions such as pathological gambling manifest in loss-chasing, where losses are misattributed to random misfortune. Despite broad clinical recognition of this link, the computational mechanisms through which uncertainty shapes learning, and through which they become disrupted in psychopathology, remain insufficiently understood. Identifying these mechanisms is critical for advancing a precise, transdiagnostic account of psychiatric dysfunction.

Modern computational neuroscience indicates that error-driven learning depends on both the prediction error (the difference between an outcome and its expectation) and the learning rate, which determines how strongly each new outcome updates subsequent beliefs ^7–10^. Statistical principles further show that the appropriate learning rate depends on two distinct sources of uncertainty: volatility and stochasticity. Volatility refers to changes in the underlying latent state itself, so that previously learned information may quickly become outdated. Stochasticity, in contrast, refers to noise in the mapping between latent state and observation, so that outcomes may be unreliable even when the environment is stable. The two sources have opposing implications for learning: volatility calls for higher learning rates so that new evidence can override outdated beliefs, whereas stochasticity calls for lower learning rates so that noisy outcomes do not displace accurate ones. The core computational challenge is therefore to correctly attribute each surprising outcome to its source ^11^.

Building on these statistical principles, we have previously developed a framework in which volatility and stochasticity are jointly inferred from experience and shown experimentally that human participants dissociate the two in both continuous and binary outcome tasks ^12,13^. The dominant paradigm in computational psychiatry, however, has continued to rely on hierarchical models in which volatility is the primary driver of learning rate modulation and stochasticity is either fixed, assumed known, or absorbed into other parameters. This paradigm has been widely used to characterize learning differences across anxiety, psychosis, autism, gambling disorders, and related conditions ^4,14–20^. Because it does not jointly infer both sources, any individual difference attributed to “volatility sensitivity” may in fact reflect a confounded mixture of volatility and stochasticity processing. How psychiatric dimensions map onto the dissociated computations, and whether the same principles generalize across reward and loss contexts, therefore remains an open question.

Our theoretical framework makes specific, testable predictions about how failures to dissociate volatility and stochasticity should produce qualitatively distinct learning phenotypes ^11^. An intact learner correctly dissociates the two sources and shows the normative pattern: increased learning under volatility, decreased learning under stochasticity (Fig. 1b, e). A stochasticity-blind learner lacks the module that attributes noise to observation and therefore misinterprets stochastic noise as evidence of environmental change, producing elevated learning under high stochasticity (Fig. 1a, d). A volatility-blind learner shows the opposite failure, misattributing genuine changes in the latent state to noise and producing reduced learning under high volatility (Fig. 1c, f). Critically, the absence of one uncertainty representation does not merely eliminate its corresponding effect on learning; it allows the remaining module to dominate, yielding a reversal in the modulation pattern that serves as a signature diagnostic for each phenotype.

**Fig. 1.**
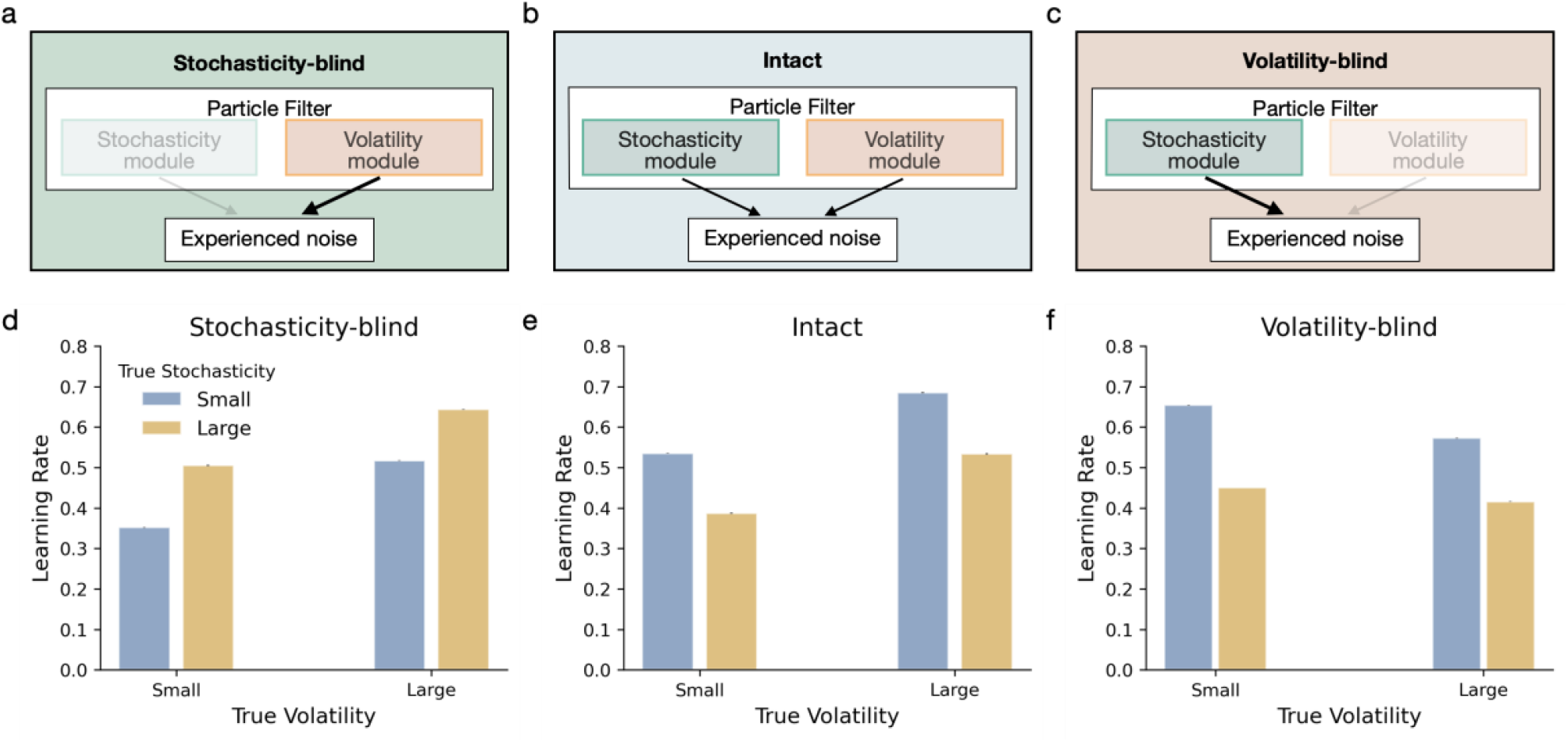
Three phenotypes of joint volatility-stochasticity inference and their reversed learning rate signatures. **a–c.** Computational architectures of the three phenotypes. The intact model (**b**) contains both stochasticity and volatility modules, which compete to explain observed noise. The stochasticity-blind (**a**) and volatility-blind (**c**) models each lack one module, so noise from the missing source is misattributed to the remaining one. **d–f**. Learning rate from simulation of 200 trials under low and high values of volatility and stochasticity. The intact model (**e**) shows the normative pattern: learning rate increases with volatility and decreases with stochasticity. Both blind models reverse this pattern on the dimension they cannot represent: the stochasticity-blind model (**d**) shows an elevated learning rate under increasing stochasticity, and the volatility-blind model (**f**) shows a reduced learning rate under increasing volatility. Error bars are standard error of the mean across simulations.

This computational architecture predicts selective rather than generalized impairments across psychopathology. Stochasticity-blindness would produce a tendency to interpret noisy outcomes as evidence of meaningful change, inflating perceived environmental shifts and driving the chronic worry, intolerance of uncertainty, self-blame, and overgeneralized avoidance that characterize anxiety and depression ^21–23^; patients with anxiety disorders show higher win-stay/lose-shift behavior than controls even when losses are due to chance ^24,25^ and benefit less from stable cue-outcome contingencies ^1,18^. Volatility-blindness would produce the opposite signature: a failure to update beliefs when the environment meaningfully shifts, consistent with the perseverative, loss-chasing behavior characteristic of pathological gambling ^6^ and with the reversal-learning and contingency-learning deficits reported in behavioral addictions ^26,27^.

We address these questions in two empirical studies. Study 1 used a continuous reward-based task that independently manipulates volatility and stochasticity, enabling model-neutral quantification of trial-by-trial updating in a large online sample (*N* > 2,600). This study characterizes individual differences in baseline learning, volatility sensitivity, and stochasticity sensitivity, and identifies subgroups corresponding to the intact, stochasticity-blind, and volatility-blind phenotypes predicted by the framework. Study 2 (*N* ≈ 700) examines whether these computational signatures generalize to binary outcomes and whether they vary with outcome valence (reward seeking vs. loss avoidance), motivated by prior work suggesting that uncertainty processing may interact with valence in anxiety ^4^. Based on the framework above, we hypothesized that anxiety and depression symptoms would be associated with elevated learning under high stochasticity (the stochasticity-blind signature), whereas behavioral-addiction symptoms would be associated with reduced learning under high volatility (the volatility-blind signature).

## Results

### Experiment 1

We first examined learning in the bird task, a prediction inference paradigm recently validated in large-scale experiments ^12^, in which volatility and stochasticity are manipulated independently in a 2 × 2 factorial design across four blocks. The task was administered to a large unselected sample recruited through Prolific (*N* = 2,532 after exclusions; see Methods and Supplementary Table 1 for demographics). Participants inferred the location of a hidden bird (the latent state) moving along a horizontal axis and positioned a bucket to catch bags of coins it dropped (Fig. 2a). Two independent sources of noise corrupted the observed bag location: volatility, the trial-by-trial change in the hidden state, and stochasticity, noise in the mapping from the hidden state to where the bag landed. The cover story framed these as the bird’s noisy movement and wind perturbations, respectively, so that participants understood the two sources as conceptually distinct. Across the four blocks, participants interacted with four different birds under four different weather conditions, with true volatility and stochasticity fixed within block but varied between blocks. Because neither value was disclosed, participants had to infer volatility and stochasticity from the observed sequence of bag positions alone in order to behave adaptively in each condition.

**Fig. 2.**
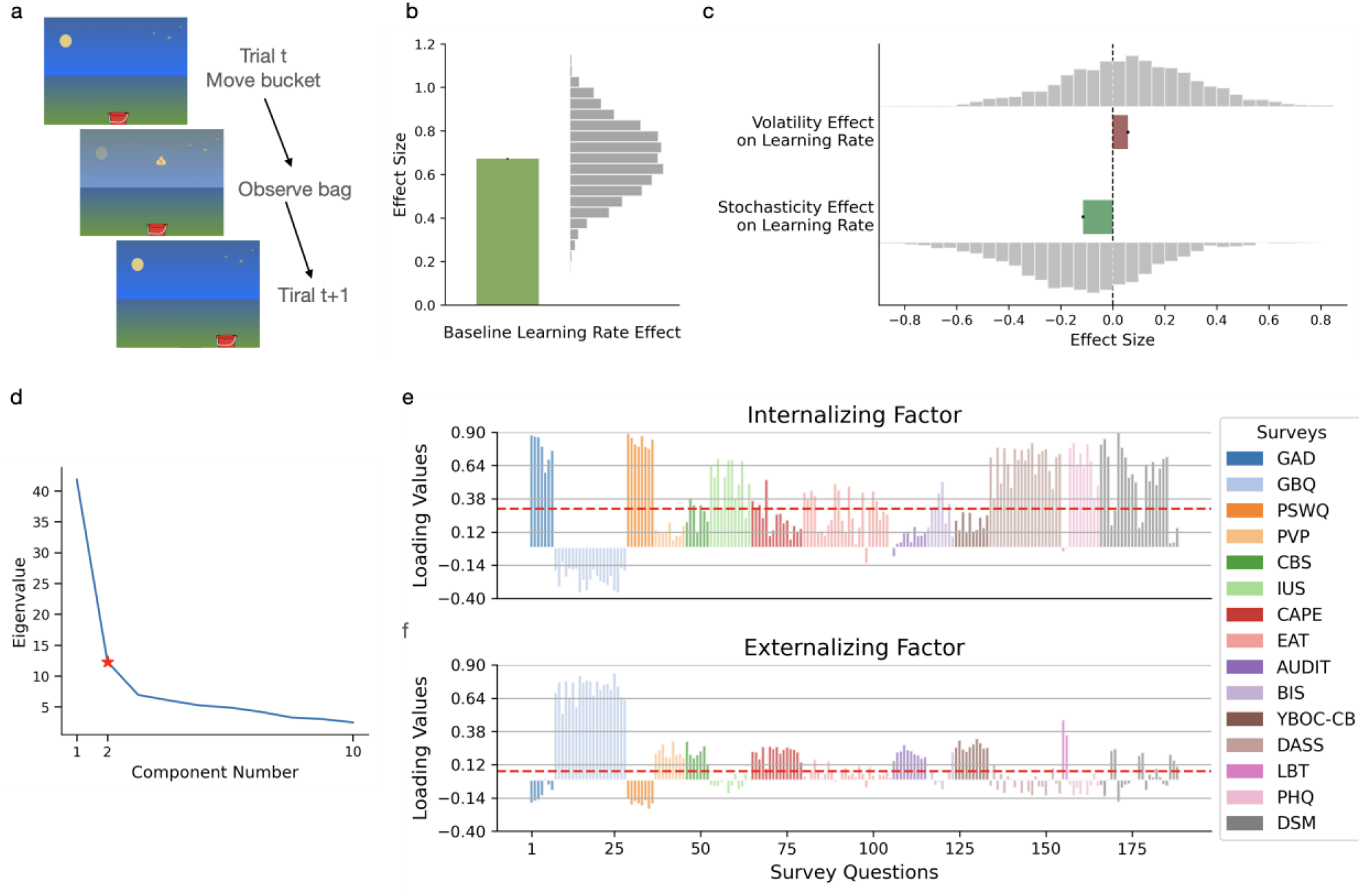
Experiment 1. **a** On every trial, participants move the bucket to catch a bag of coins dropped by an invisible bird. Participants cannot move the bucket when the bag appears and the screen turns opaque. The task has four blocks with a 2 × 2 factorial design, manipulating both true volatility and true stochasticity. **b–c** Empirical learning rate, estimated per block from a model-neutral regression of belief updates on prediction errors. **b** Baseline learning rate (block-averaged). **c** Main effects of volatility and stochasticity on learning rate, each computed as the difference between corresponding high and low blocks. Mean and standard error of the mean are plotted. **d** Scree plot illustrating eigenvalues derived from exploratory factor analysis from Experiment 1, indicating two significant latent factors. **e-f** Factor loading values for the two identified factors across all questionnaire items (188 items total). Questionnaire sources are differentiated by color.

This design enabled a model-agnostic analysis of learning rate adjustment under uncertainty. We defined learning rate as the ratio of belief updating to prediction error: the update was the trial-by-trial change in bucket position, and the prediction error was the difference between the observed bag location and the current bucket position. For each participant, we quantified block-wise learning rates by regressing trial-by-trial updates onto prediction errors, controlling for block-specific differences unrelated to prediction error (see Methods). Across the sample, the two uncertainty sources had opposing effects, as predicted (Fig. 2c): participants increased their learning rate in high-volatility blocks (*t*(2531) = 10.72, *P* < 0.001) and decreased it in high-stochasticity blocks (*t*(2531) = −22.06, *P* < 0.001), with no significant interaction (*t*(2531) = −1.04, *P* = 0.298; Supplementary Table 2). These findings replicate Piray & Daw ^12^ and confirm that, at the population level, participants appropriately dissociated the two sources of uncertainty. Substantial heterogeneity across individuals in baseline learning, volatility sensitivity, and stochasticity sensitivity (Fig. 2b–c) motivates the analyses that follow, which link these learning profiles to latent psychiatric dimensions.

After the bird task, participants completed a comprehensive battery of self-report questionnaires assessing a broad range of psychiatric symptoms, including anxiety, worry, intolerance of uncertainty, depression, impulsivity, gambling, alcohol use, compulsive behavior, problem video-game playing, disordered eating, and psychotic-like experiences (see Methods for the full list of instruments). To identify latent dimensions of psychopathology, we conducted exploratory factor analysis on the full questionnaire set. The scree plot showed a marked drop after the second eigenvalue, supporting a two-factor solution (Fig. 2d). The first factor loaded most strongly on measures of anxiety and depression (GAD-7, PSWQ-8, IUS-12, DASS-21, PHQ-9), and to lesser extent on specific questions of eating disorders and impulsivity (EAT-26, BIS-11; Fig. 2e). The second factor was driven primarily by gambling-related measures (GBQ-21, LBT), with smaller loadings from instruments assessing problem video-game playing (PVP-9), compulsive buying (CBS-7, YBOC-CB-10), psychotic-like experiences (CAPE-15), and alcohol use (AUDIT) (Fig. 2f). These loading patterns are broadly consistent with transdiagnostic Internalizing and Externalizing dimensions, though we note that the symptom coverage of our battery is restricted to these specific domains rather than spanning the full range of either dimension as defined in HiTOP-style frameworks (see Methods, Factor analysis). For brevity, we refer to these two factors as Internalizing and Externalizing in the remainder of the paper and use participant-level factor scores in subsequent analyses.

Motivated by the theoretical framework, we next asked whether individual differences in learning rate effects could identify the three predicted computational phenotypes. Each blind variant produces a characteristic reversal: a stochasticity-blind learner shows a positive stochasticity effect, because the unopposed volatility module attributes all surprise to environmental change; symmetrically, a volatility-blind learner shows a negative volatility effect, because the unopposed stochasticity module attributes all surprise to noise. The present study was designed with a sufficiently large sample to isolate these relatively extreme phenotypes and test whether they map onto distinct psychiatric profiles.

To detect these signatures empirically, we defined two critical values estimated from an independent, previously published sample of the same task (*N* = 643): *c*_*v*_, the average volatility effect (a positive value), and *c*_*s*_, the average stochasticity effect (a negative value). Using these thresholds, we classified participants into three groups. Intact participants (*N* = 526) showed a volatility effect greater than *c*_*v*_ and a stochasticity effect more negative than *c*_*s*_, exhibiting clear sensitivity to both sources of uncertainty in the expected directions (Fig. 3b). Stochasticity-blind participants (*N* = 349) showed a stochasticity effect greater than *c*_*v*_ (positive, the wrong direction) together with a volatility effect also greater than *c*_*v*_ (preserved volatility sensitivity), exhibiting reversed responses to stochasticity alongside intact responses to volatility (Fig. 3a). Symmetrically, volatility-blind participants (*N* = 315) showed a volatility effect less than *c*_*s*_ (negative, the wrong direction) together with a stochasticity effect also less than *c*_*s*_ (preserved stochasticity sensitivity), exhibiting reversed responses to volatility alongside intact responses to stochasticity (Fig. 3c). This dual-criterion structure ensures that group membership reflects reliable computational signatures rather than measurement noise. Critically, both blind phenotypes require intact sensitivity to the other source of uncertainty, capturing selective deficits in attribution rather than global task disengagement or generally elevated or diminished learning.

**Fig. 3.**
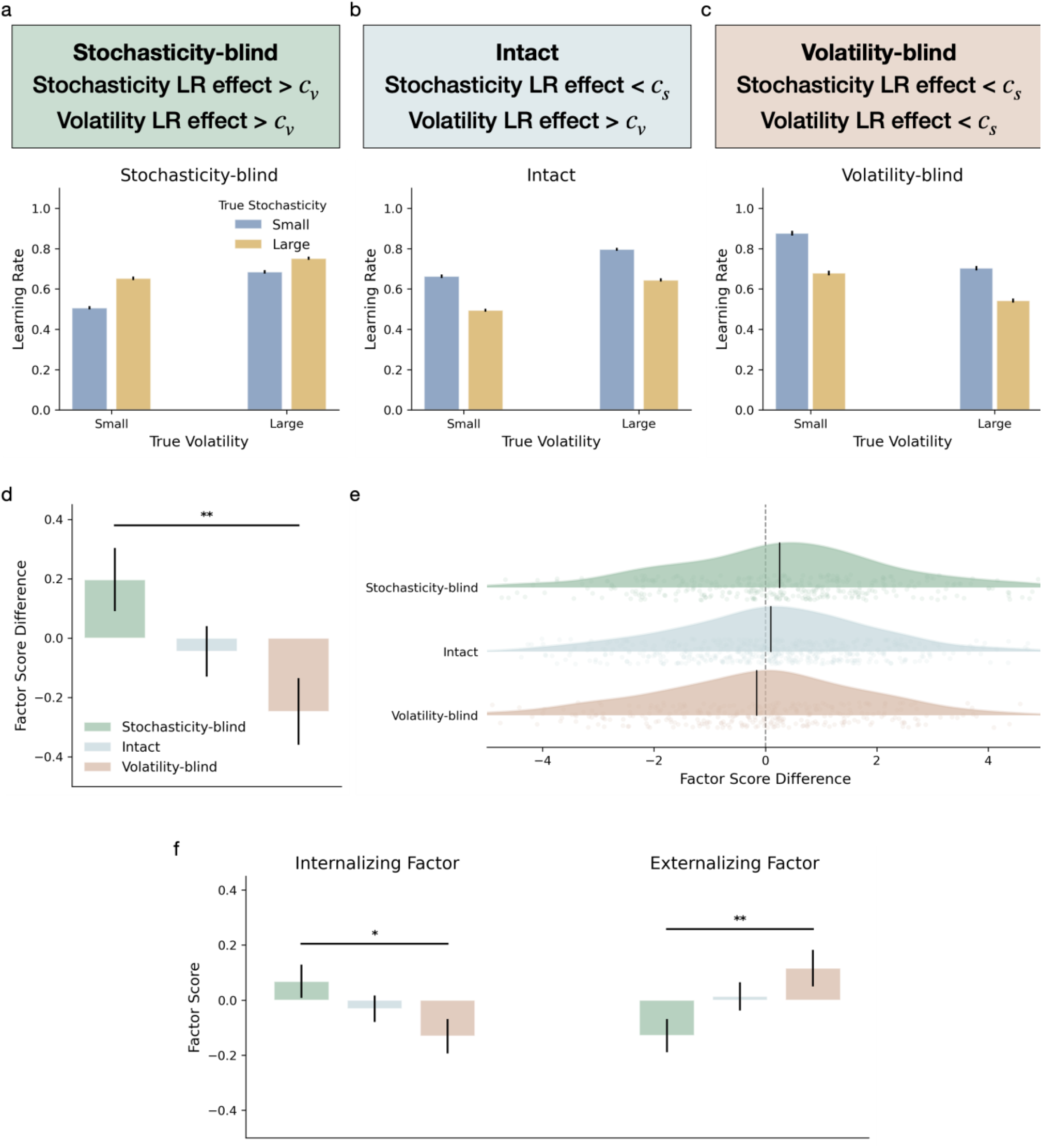
Three computational phenotypes show a double dissociation with transdiagnostic psychiatric dimensions. **a-c** Participants in Experiment 1 were classified according to their model-neutral stochasticity and volatility effect on learning rate relative to critical values *c*_*s*_ and *c*_*v*_ , defined as the mean stochasticity and volatility effects from an independent previously published large-scale sample^12^. The critical values were defined as the pilot-sample mean stochasticity effect (*c*_*s*_) and mean volatility effect (*c*_*v*_). Stochasticity-blind participants (*n* = 349; **a**) were defined as those with stochasticity effect and volatility effect above the *c*_*v*_, indicating exaggerated updating under stochastic noise despite preserved sensitivity to volatility. Intact participants (n = 526; **b**) had volatility effects above *c*_*v*_ and stochasticity effects below *c*_*s*_, consistent with adaptive dissociation of the two uncertainty sources. Volatility-blind participants (n = 315; **c**) had both effects below *c*_*s*_, indicating inverse sensitivity to volatility despite preserved sensitivity to stochasticity. Mean and standard error of the mean are plotted. **d** Difference (Internalizing − Externalizing) score for each phenotype. Stochasticity-blind participants showed positive differences (higher Internalizing); volatility-blind participants showed negative differences (higher Externalizing). **e** Distribution of the factor score difference across the three phenotype groups, shown as half-violin density plots with individual participant data overlaid. Vertical black lines indicate group means, and the dashed vertical line marks zero. For illustration purpose here, outliers were excluded using the 1.5 × IQR criterion within each group. **f** Factor scores for each computational phenotype. Asterisks indicate significant difference between blind groups, with *P* < 0.01 (∗∗), *P* < 0.05 (∗).

These computational phenotypes mapped onto distinct psychiatric profiles, consistent with the theoretical prediction of a double dissociation between blind variants and symptom dimensions (Fig. 3d-f). A mixed-design ANOVA with Groups (intact, stochasticity-blind, volatility-blind) as a between-subjects factor and Dimensions (Internalizing, Externalizing) as a within-subjects factor revealed a significant Group × Dimension interaction (*F*(2,1187) = 4.23, *P* = 0.015; Fig. 3d), indicating that the relative balance of Internalizing and Externalizing scores differed across the three groups. This interaction was driven primarily by a sharper contrast between the two blind groups: stochasticity-blind participants showed relatively higher Internalizing than Externalizing scores, whereas volatility-blind participants showed the opposite pattern (*F*(1,662) = 8.30, *P* = 0.004; Fig. 3e).

Examining the two dimensions separately confirmed the same pattern (Fig. 3f). On the Internalizing dimension, the stochasticity-blind group scored highest, the intact group intermediate, and the volatility-blind group lowest, with a significant difference between the two blind groups (*F*(1,662) = 5.23, *P* = 0.023); the main effect of Group was non-significant (*F*(2,1187) = 2.65, *P* = 0.071). On the Externalizing dimension, the volatility-blind group scored highest, with a significant main effect of Group (*F*(2,1187) = 3.75, *P* = 0.024) again driven by the contrast between the two blind groups (*F*(1,662) = 7.48, *P* = 0.006; Supplementary Table 3). Together, these results indicate a selective mapping: failure to attribute stochasticity is associated with anxiety- and depression-type symptoms, whereas failure to attribute volatility is associated with behavioral-addiction symptoms.

To complement the categorical grouping analysis with a population-level view, we next examined how learning rate and its sensitivities to volatility and stochasticity relate to psychiatric dimensions across the whole sample (Supplementary Fig.1; Supplementary Table 4). This analysis addresses a methodological limitation of prior work: in designs that manipulate only volatility, an elevated baseline learning rate cannot be cleanly distinguished from elevated volatility sensitivity. Individuals with high baseline learning rate (for example, anxious participants) approach the learning rate ceiling of 1, which artificially suppresses their measurable volatility slope and conflates baseline differences with apparent differences in volatility sensitivity. The factorial manipulation of stochasticity in our design mitigates this confound: high-stochasticity blocks pull learning rate below ceiling, providing conditions in which volatility-driven changes can be observed without saturation. In addition, each task-derived quantity is computed from an orthogonal contrast across the 2×2 factorial cells, so baseline learning rate, volatility sensitivity, and stochasticity sensitivity are unconfounded by design. We regressed each task-derived computational quantity onto the two psychiatric factor scores (Internalizing, Externalizing) using general linear models (see Methods).

This dimensional analysis converged with the phenotype-based results while providing a more continuous characterization of the same dissociation. Higher Internalizing scores predicted greater baseline learning (*t*(2529) = 3.66, *P* < 0.001), whereas baseline was not associated with Externalizing (*P* = 0.964). The volatility effect, by contrast, was selectively associated with Externalizing: higher Externalizing scores predicted a reduced volatility effect (*t*(2529) = −3.29, *P* = 0.001), whereas the volatility effect was not associated with Internalizing (*P* = 0.487). The stochasticity effect was not associated with either dimension (Internalizing: *P* = 0.434; Externalizing: *P* = 0.236). Thus, although presented here as supportive evidence, the whole-sample dimensional analysis helps clarify that Internalizing and Externalizing dimensions were related to separable components of uncertainty-guided learning rather than to a single undifferentiated change in updating. .

### Experiment 2

Experiment 1 established a double dissociation between computationally-defined learner phenotypes and psychiatric dimensions: stochasticity-blind learners scored higher on Internalizing, while volatility-blind learners scored higher on Externalizing. At the population level, these findings were complemented by selective associations of Internalizing with elevated baseline learning and Externalizing with reduced volatility sensitivity. We next asked whether these population-level associations generalize to binary learning environments and whether they differ between reward-based and loss-based versions, motivated by prior work suggesting that uncertainty processing may interact with valence in anxiety ^4^. The continuous bird task does not lend itself to a clean reward-versus-loss comparison, so we used a pair of recently validated binary tasks ^13^ that retain the factorial manipulation of volatility and stochasticity and differ only in outcome valence. Unlike continuous feedback, binary feedback offers clear-cut evaluations of success or failure, which may exacerbate maladaptive learning patterns and may partly explain why people learn differently from positive versus negative prediction errors ^28–30^.

The two binary tasks shared a 2 × 2 factorial design in which true volatility and true stochasticity were manipulated independently across four blocks of 40 trials, with parameters fixed within each block and not disclosed to participants (*N* = 723, after exclusions; see Methods). In the reward-based task, participants chose between two sides of a beach on each trial, attempting to find treasure left by a hidden sea lion whose position they had to infer from outcomes (Fig. 4a). Participants were instructed that the sea lion typically remained on the same side across trials but could suddenly switch (corresponding to volatility), and ocean waves could independently displace where the treasure appeared (corresponding to stochasticity). The loss-based task was structurally identical but used a hidden turtle: choosing the side the turtle had visited could result in a jellyfish sting and lost points, so participants were instructed to avoid that side (Fig. 4b). In each task, participants completed four blocks, each featuring a different sea lion (in the reward task) or turtle (in the loss task) on a different beach, and were explicitly instructed that the animal’s switching and the wave perturbations were independent sources of noise.

**Fig. 4.**
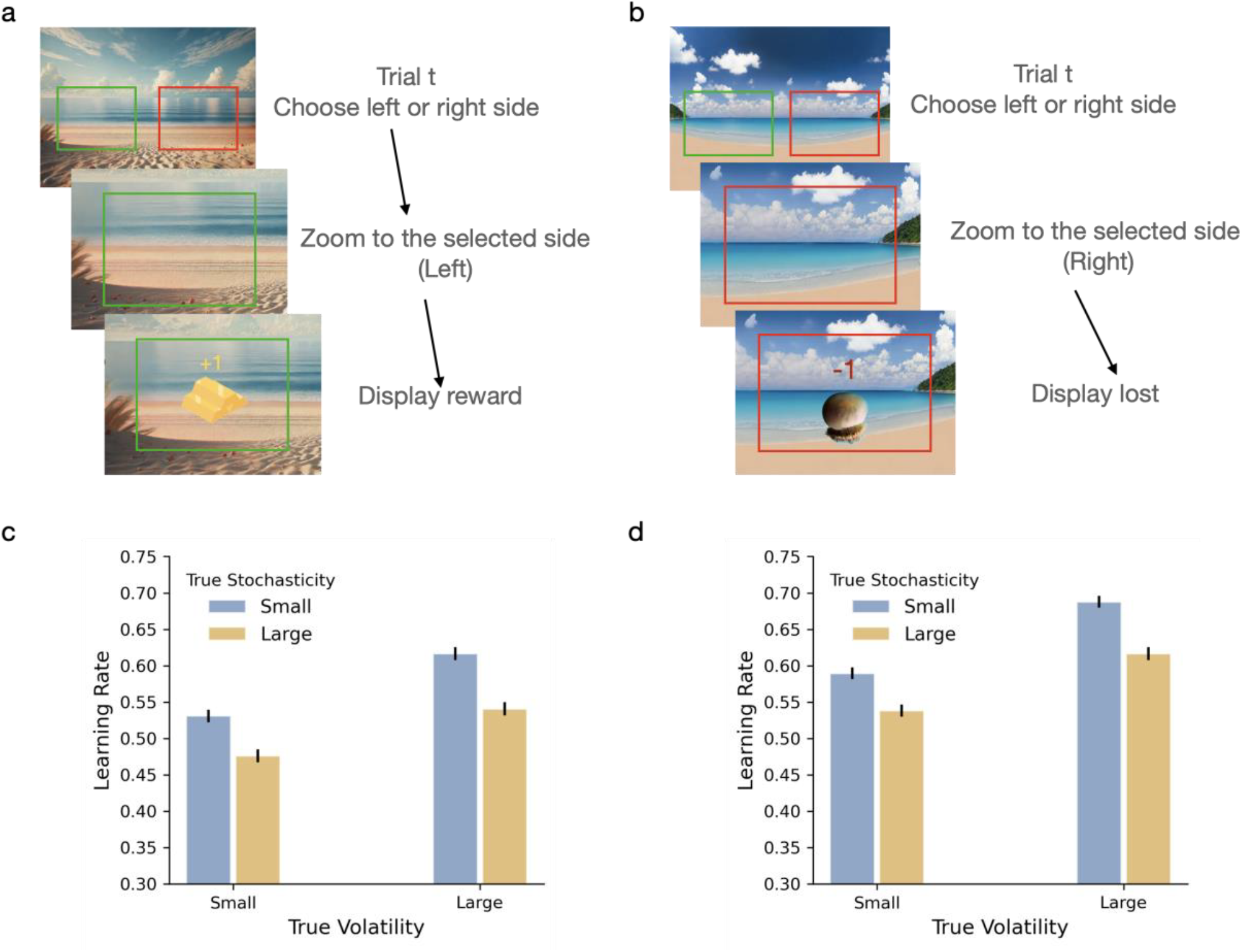
Binary tasks design and learning rate in Experiment 2. a–b, On each trial, participants choose one side of the beach using the left or right arrow key; the screen then zooms in to display the outcome. The two tasks share an identical 2 × 2 factorial structure similar to the bird task, and differ only in outcome valence: in the reward-based task (**a**), participants find treasures brought by a hidden sea lion (reward or no reward); in the loss-based task (**b**), they avoid jellyfish brought by a hidden turtle (point loss or no loss). The chosen sides in the illustrations (left, green box in a; right, red box in b) are not displayed in the actual task. c–d, Mean learning rate per block from CBF-HMM fits (N = 723) for the reward-based (c) and loss-based (d) tasks. Both tasks show the normative pattern: learning rate increases under higher volatility and decreases under higher stochasticity. Error bars indicate standard error of the mean.

In the binary tasks, the model-neutral regression used in Experiment 1 cannot be applied directly, because both predictions and outcomes are binary. We instead analyzed the binary tasks using a Hidden Markov Model (HMM) framework, which provides the normative inference structure for binary hidden states and binary observations and supports joint inference of volatility and stochasticity ^13^. Specifically, we used a categorical Bayes filter (CBF) ^31^ which implements this inference in a way that is robust for individual-differences analyses. We fit the model separately to each participant’s data, yielding trial-by-trial predicted probabilities. Using the best-fit parameters for each participant, we generated trial-wise predictions and derived block-wise estimates of learning rate for both binary tasks (see Methods, Model fitting).

The model-based learning rates revealed that the adaptive patterns identified in the continuous bird task generalize to binary environments. Across both reward- and loss-based tasks, participants successfully differentiated the two uncertainty sources, increasing their learning rate under high volatility and decreasing it under high stochasticity (Fig. 4c, d). In the reward-based task, true volatility had a significant positive effect on learning rate (*t*(722) = 19.93, *P* < 0.001), whereas true stochasticity had a significant negative effect (*t*(722) = −27.36, *P* < 0.001), along with a significant baseline learning effect (*t*(722) = 63.89, *P* < 0.001) (Supplementary Table 5, Supplementary Fig. 2a, b). In the loss-based task, the same pattern was observed, with a significant positive effect of true volatility (*t*(722) = 22.85, *P* < 0.001) and a significant negative effect of true stochasticity (*t*(722) = −25.33, *P* < 0.001), along with a significant baseline learning effect (*t*(722) = 76.16, *P* < 0.001) (Supplementary Table 6, Supplementary Fig. 2c, d). These results replicate our recent work ^13^, and establish that the core learning patterns identified in the bird task generalize to binary decision settings and remain evident across both reward-seeking and punishment-avoidance contexts.

Following the binary tasks, participants completed a subset of the surveys used in Experiment 1, selected to represent the Internalizing and Externalizing dimensions while minimizing participant burden. To ensure a consistent psychiatric framework across the reward and loss contexts, we conducted a factor analysis on the combined questionnaire data from both sessions (see Methods). This analysis recovered the two primary factors corresponding to the Internalizing and Externalizing dimensions (Supplementary Fig. 3). The resulting factor scores were used in all subsequent analyses to relate learning behavior across both valence contexts to a common psychiatric space.

To examine how learning depended on outcome valence, we decomposed each participant’s learning-rate components into effects shared across reward and loss contexts and effects specific to outcome valence. For each task-driven computational quantity, i.e., baseline learning rate, stochasticity effect, volatility effect, we computed the average value across both tasks (capturing the participant’s general tendency under binary uncertainty) and a valence contrast (the within-subject difference between loss and reward; Fig. 5a, b). Group-level valence effects on each component are reported in Supplementary Table 7. We next used linear regression models to test how each psychiatric dimension related to the task-averaged components and the valence contrasts (Fig. 5c; see Methods; Supplementary Table 8, Supplementary Fig. 4).

**Fig. 5.**
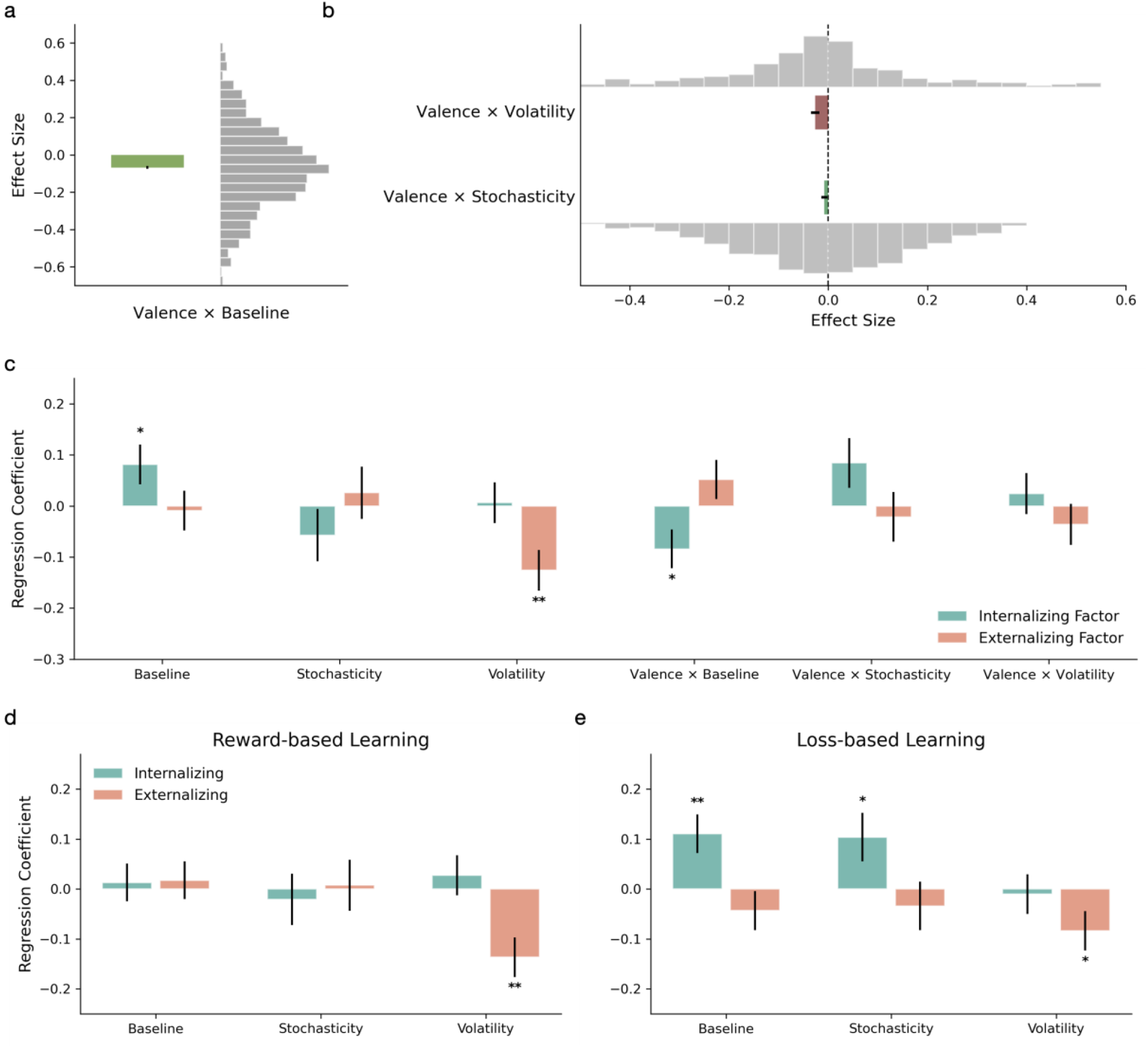
Shared and valence-dependent associations between binary tasks learning effects and psychiatric factors. **a** Distribution of the Valence × Baseline term (within-subject difference in baseline learning rate between the reward- and loss-based tasks). **b** Distributions of the Valence × Volatility and Valence × Stochasticity terms (within-subject differences between the two tasks in the volatility and stochasticity effects, respectively). In **a–b**, colored bars indicate mean effects and gray histograms show participant-level distributions. **c** Regression coefficients from regression models relating binary-task learning rate effects to psychiatric dimensions. Predictors included the baseline learning rate (Baseline), stochasticity effect (Stochasticity), and volatility effect (Volatility) across the two tasks, together with the corresponding valence interaction terms quantifying within-subject differences between the two tasks. Coefficients are shown separately for the Internalizing and Externalizing factors. **d–e**, Task-specific GLM coefficients for the reward-based task (**d**) and loss-based task (**e**); predictors included baseline learning rate, stochasticity effect, and volatility effect. Error bars indicate standard errors. Asterisks indicate statistical significance, with *P* < 0.005 (∗∗) and *P* < 0.05 (∗).

For the Internalizing dimension, the task-averaged baseline learning rate showed a significant positive association (*t*(714) = 2.09, *P* = 0.037): higher Internalizing scores were linked to stronger overall updating across both binary tasks, replicating the Experiment 1 baseline-Internalizing finding. Neither the task-averaged stochasticity effect (P = 0.264) nor the task-averaged volatility effect (*P* = 0.871) was associated with Internalizing, also consistent with Experiment 1. In addition, the valence contrast for baseline learning was negatively associated with Internalizing (*t*(714) = −2.21, *P* = 0.028); given that the group-level valence contrast for baseline was negative (reward < loss), this indicates that the Internalizing-baseline association was stronger in the loss-based task than in the reward-based task. Higher Internalizing scores were thus associated with elevated baseline updating in both tasks, with a larger elevation under loss.

For the Externalizing dimension, only the task-averaged volatility effect showed a significant association (*t*(714) = −3.16, *P* = 0.002), indicating that participants with higher Externalizing scores showed weaker volatility-driven increases in learning rate across both binary tasks. No other task-averaged effect or valence contrast was significantly associated with Externalizing. This pattern replicates the Experiment 1 finding that Externalizing is selectively linked to reduced volatility sensitivity and indicates that this signature generalizes across reward and loss contexts without valence-specific modulation. Reduced volatility sensitivity thus emerges as a stable computational signature of the Externalizing dimension, present in both continuous and binary settings and invariant to outcome valence.

To localize these effects across reward and loss contexts, we also ran task-specific regressions for each binary task separately (Fig. 5d, e; Supplementary Tables 9-12). The Externalizing-volatility association was present in both the reward-based task (*t*(718) = −3.44, *P* = 0.001) and the loss-based task (*t*(718) = −2.11, *P* = 0.035), confirming that this signature is consistent across valence. The Internalizing-baseline association, in contrast, was significant only in the loss-based task (*t*(718) = 2.85, *P* = 0.005), consistent with the loss-amplified valence interaction reported above. In the same task, Internalizing also showed a significant positive association with the stochasticity effect (*t*(718) = 2.15, *P* = 0.032), indicating reduced dampening of learning under stochastic noise specifically in the aversive context.

## Discussion

Distinct sources of uncertainty can shape learning, and individual differences in these computations is related to latent psychiatric dimensions ^1–6^. In all three tasks, participants showed the normative pattern predicted by joint-inference theories of uncertainty ^11–13^: learning rate increased under high volatility and decreased under high stochasticity. Against this common behavioral background, a theory-guided categorical analysis in Experiment 1 identified three distinct computational phenotypes predicted by the framework: intact, stochasticity-blind, and volatility-blind learners. Crucially, the phenotypes mapped onto distinct psychiatric profiles in a double-dissociation pattern: stochasticity-blind learners scored higher on Internalizing, while volatility-blind learners scored higher on Externalizing. This finding provides direct empirical support for a selective rather than generalized account of computational deficits in psychiatry, in which distinct failure modes of uncertainty attribution correspond to distinct dimensions of psychopathology.

Dimensional analyses across both experiments converged with and extended the categorical findings. The Internalizing dimension was selectively associated with elevated baseline learning rate, while the Externalizing dimension was selectively associated with reduced volatility sensitivity, and these two associations generalized from continuous outcomes (Experiment 1) to binary outcomes (Experiment 2). In the matched binary reward and loss tasks, the Internalizing-baseline association was further amplified in the loss context, suggesting that aversive contexts intensify Internalizing-related over-updating. By contrast, the Externalizing-volatility association was invariant to valence, indicating a stable signature of reduced responsiveness to environmental change across reward and loss. Together, these findings support the view that maladaptive learning under uncertainty is better understood as a set of dissociable computational phenotypes than as a single generalized impairment.

Within this framework, the three learner phenotypes are not arbitrary clusters but specific predictions of the joint-inference theory ^11–13^. The intact learner correctly attributes surprise to either volatility or stochasticity, adjusting learning rate adaptively. Each blind phenotype reflects a failure of this attribution: the stochasticity-blind learner treats stochasticity as volatility and over-updates, while the volatility-blind learner treats volatility as stochasticity and under-updates. Our earlier work has shown that this attribution step is the computationally most demanding part of joint inference, with the corresponding adjustment of learning rate following in a relatively straightforward way once attribution is made ^11–13^. The step is also behaviorally pivotal, since volatility and stochasticity demand opposite adjustments of learning rate: misattributing one for the other reverses the direction of updating rather than scaling it. Because attribution is both inferentially difficult and behaviorally consequential in this way, it is the natural locus at which systematic biases and individual differences in psychiatric symptomatology should emerge. The defining behavioral signature of these biases is a selective reversal of learning rate sensitivity, in which the response to the source the learner cannot perceive takes the wrong sign. This reversal distinguishes the phenotype framework from accounts that treat learning rate as a single continuous parameter, since no such account predicts that learning rate sensitivity can reverse direction. The mapping we observe, with stochasticity-blindness enriched for Internalizing and volatility-blindness enriched for Externalizing, therefore identifies not two graded associations but two qualitatively distinct ways in which uncertainty processing can go awry.

The categorical analyses confirmed our hypothesis and prior modeling work: individuals classified as stochasticity-blind were significantly enriched for Internalizing, reflecting a selective failure to dampen learning rate under high stochasticity. One cognitive process that could contribute to this pattern is self-blame: an anxious individual who experiences a negative outcome that might in fact be due to chance may attribute it to themselves and change course abruptly, rather than dampening the update as a chance origin would warrant. The dimensional analyses across the entire population revealed an additional finding not directly predicted by the framework: Internalizing was selectively associated with elevated baseline updating across both experiments. The Internalizing-baseline finding is consistent with broader observations of elevated updating in anxiety and depression, though the precise computational locus has been debated. Earlier studies emphasized altered volatility sensitivity ^1,4^, but more recent better-powered work has produced inconsistent results, with some larger studies failing to replicate the effect ^32^. One methodological consideration is that volatility-only paradigms have limited dynamic range for estimating volatility-related learning-rate changes, particularly when participants have elevated baseline updating that pushes learning rate toward ceiling during volatile blocks; this can make any volatility-related individual differences difficult to detect, regardless of whether they exist. The factorial design used here mitigates this constraint and identifies the Internalizing association at baseline rather than at volatility sensitivity. In addition, our analytic approach used transdiagnostic latent factors derived from a broad symptom battery, whereas studies relying on clinically-oriented self-report scales may have reduced sensitivity to the dimensional variation captured here. The factorial design and dimensional symptom measurement used here mitigate these constraints and identify the Internalizing association at baseline rather than at volatility sensitivity.

The valence finding is particularly informative because prior work on reward versus loss learning in anxiety has not provided a consistent computational setup. Some studies have emphasized greater sensitivity to punishment or aversive feedback ^4,28^, whereas others have found weak or null effects of valence on learning rate adaptation itself ^18^. In many cases, reward and punishment conditions were not structurally matched, or the models used did not distinguish baseline updating from volatility-related and stochasticity-related components. Our design addressed this gap directly by comparing reward- and loss-based learning under the same 2 × 2 factorial design. Within this framework, Internalizing was associated with a stronger loss-over-reward shift in baseline updating, suggesting that valence selectively amplifies general responsiveness to evidence in this dimension. A complementary exploratory finding in the loss-based task showed that Internalizing was also positively associated with the stochasticity effect itself, suggesting reduced normative dampening of learning under stochastic noise specifically under loss. This single significant test should be interpreted with caution, but it points to the same stochasticity-related component identified by the categorical stochasticity-blind classification. The valence amplification in Internalizing thus operates on baseline updating and stochasticity-related computations, leaving volatility sensitivity unaffected. These findings position Internalizing as showing both a predicted categorical stochasticity-blindness pattern and an additional dimensional signature of elevated baseline updating, both becoming more pronounced under aversive conditions.

The Externalizing dimension in the present study was derived from a symptom set that primarily captures behavioral addiction and compulsivity, and is therefore narrower than the broader Externalizing construct used in some psychometric frameworks, which also includes conduct and antisocial dimensions. Within this scope, both the categorical and dimensional analyses identified the same computational locus: volatility sensitivity. Categorically, individuals classified as volatility-blind were significantly enriched for Externalizing, reflecting a selective failure to raise learning rate when environmental change requires it. Dimensionally, higher Externalizing scores were associated with dampened increases in learning rate following contingency changes, across both Experiment 1 and Experiment 2. The two analyses thus provide convergent evidence at different levels of severity for the same Externalizing-volatility association anticipated by joint-inference theory. A learner with reduced volatility sensitivity may continue relying on outdated beliefs even when the underlying contingencies have changed, sustaining maladaptive choices. This characterization is consistent with broader work linking impulsivity, compulsivity, and addiction-related behavior to impaired behavioral flexibility and difficulty adapting to contingency reversals ^33^, and resonates with computational theories of compulsive and addictive behavior in which salient losses or contingency changes fail to produce appropriate behavioral correction, contributing to persistence and loss chasing ^34– 36^. The Externalizing-volatility association did not vary with valence; it appeared in both reward-based and loss-based binary tasks, implying a more general failure to register and respond to environmental change rather than a selective bias toward one outcome type.

Several methodological features of the present work warrant emphasis. Our computational analyses combine Categorical Bayes Filter ^31^ with HMMs ^13^, a variant of joint-inference modeling developed specifically for robust individual-differences analyses. The CBF yields stable trial-wise estimates of learning rate and parameter estimates well suited to the within-participant regression analyses reported here. Categorical phenotype classifications used thresholds independently derived from a previously published large-scale dataset (N = 643) ^12^ rather than fitted to the present data, ensuring that the phenotype-symptom associations are not artifacts of within-sample threshold tuning. The Experiment 1 sample size (N > 2,500) was set to ensure adequate categorical classification power, given that the strict threshold-based criteria classify approximately 45% of participants into one of the three phenotype groups; this yielded several hundred participants per group and sufficient statistical power for between-group comparisons. Combined with the Experiment 2 sample (N = 723), factorial manipulation of volatility and stochasticity across three tasks spanning continuous and binary outcomes and reward and loss valences, and transdiagnostic dimensional symptom measurement, the methodological framework provides the resolution needed to detect both categorical phenotypic patterns and dimensional component-specific associations.

Limitations of the present work should also be acknowledged. The data were collected online from a general population sample with self-report symptom measurement and without structured clinical interviews. While online attention and response quality were addressed with extensive comprehension and attention checks ^37^, the present findings should be interpreted as evidence for transdiagnostic dimensional variation rather than as disorder-specific biomarkers, and replication in clinically diagnosed samples remains an important next step. The Externalizing factor analyzed here primarily captures behavioral addiction and compulsivity and does not adequately represent the conduct, antisocial, or substance-use dimensions of broader Externalizing constructs; the Externalizing-volatility findings should therefore be regarded as specific to the symptom domains we measured. Across both experiments, stochasticity-related psychiatric associations were weaker and less consistent than those involving baseline updating or volatility sensitivity, which may indicate that individual differences in stochasticity processing are smaller or more difficult to detect behaviorally, particularly outside paradigms designed to amplify stochasticity-related inference. In Experiment 2, the reward-based and loss-based binary tasks were administered on separate days in a fixed order across participants; although within-session fatigue and carryover are not concerns under this design, residual order effects on cross-task comparisons cannot be entirely excluded.

While we focused on the learning rate consequences of volatility and stochasticity inference, the dissociation has implications for a broader set of uncertainty-sensitive computations that are themselves disrupted in psychopathology. Exploration is one example: optimal exploration policies depend on whether observed variance reflects volatility or stochasticity ^38^, and the reduced directed exploration documented in anxiety ^39^ could reflect a misattribution of variance to one source rather than the other, with downstream consequences for the avoidance behaviors that characterize the disorder. Planning is another: just as volatility and stochasticity have opposing implications for the learning rate, they should also modulate planning in distinct ways, with volatility favoring shorter planning horizons and more frequent replanning, and stochasticity favoring more cautious deliberation over noisy evidence. Theories integrating inference and planning ^40^, such as linear reinforcement learning ^41–44^, provides a natural formalism in which both noise sources can be incorporated directly into the planning machinery, and systematic biases in their inference may help explain decision-making deficits documented in anxiety and addictive disorders. Extending the joint-inference framework to these processes and characterizing how individual differences in volatility and stochasticity inference shape them, is a natural next step for relating the full breadth of decision-making impairments to transdiagnostic dimensions of psychopathology.

Future work should test these computational phenotypes in clinically diagnosed samples, examine their longitudinal stability over time and across treatment, and combine behavior with neural or physiological measures to characterize the neural mechanisms underlying the dissociation. Advancing this area more broadly requires simultaneous manipulation of both volatility and stochasticity, and modeling frameworks that, rather than holding either factor as given, focus on the central computational problem at hand: inferring what causes the experienced noise. Uncertainty has long been central to accounts of psychiatric variation; perhaps more central still is the inference individuals draw about what is causing it.

## Methods

### Experimental Procedure

The study was approved by the Institutional Review Board at the University of Southern California (“Learning and decision making under uncertainty”; Protocol Number: UP-23-00359). Participants were recruited online via the Prolific platform and provided informed consent prior to beginning the experiment. Experimental tasks were implemented in JavaScript and jsPsych ^45^ and deployed using the NivTurk platform ^46^.

Eligibility was limited to USA residents above the age of 18 years-old and are fluent in English. Participants first read instructions, completed step-by-step practice trials to familiarize themselves with the task, and then completed a comprehension quiz to ensure understanding. Those who failed the quiz twice were not allowed to proceed. Following the task, participants completed an extensive battery of self-report questionnaires (see Methods, Psychological questionnaires for more detail), additional cognitive tasks not reported here, and demographic surveys. In Experiment 1, we recruited *N* = 2689 participants. After applying the exclusion criteria, *N* = 2532 participant data were used for data analysis. In Experiment 2 reward-based task, we invited participants from the Experiment 1 pool to complete the binary tasks. From those who completed the reward-based task, *N* = 723 participants completed the loss-based task and remained for data analysis after applying exclusion criteria. All participant data used in analysis passed the comprehension check and met quality control criteria, including consistent engagement with the task (i.e., not leaving the interface idle for extended periods).

### Exclusion criteria

Inattentive responding on questionnaire items may lead to artificial correlations between task behavior and symptom measures ^37^. To identify and control for inattentive participants, we added thirteen infrequency items (e.g., “I buy things for my future alien friend.”) into questionnaires. These questions were designed to have certain correct answers. We also included two command questions (e.g., “Degree of attention over time. Select ‘I experience this rarely’ for this question.”), where correct answer is embedded in the question itself. Participants who failed more than two of the infrequency items or at least one command question were excluded from the data analysis.

### Power and sample size

Sample sizes for both experiments were determined a priori. For Experiment 1, we expected approximately 15% of participants to be classified into each of the two blind phenotype groups based on previous data ^12^. A target sample of approximately 2,500 participants therefore yields ∼350 participants per blind group, providing 80% power to detect a small effect (Cohen’s f = 0.1) and 99% power to detect a medium effect (f = 0.25) in between-group comparisons at *α* = 0.05. For Experiment 2, the target sample of approximately 750 participants was chosen to provide about 80% power for detecting small effects (standardized *β* = 0.1) at *α* = 0.05.

### Learning rate

In Experiment 1, we used participants’ choices in a model-neutral analysis to compute the per-block learning rate as the ratio of belief update to prediction error in that block. The belief update on trial *t* was defined as the change in bucket position between consecutive trials:

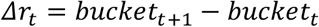

The prediction error was defined as the difference between the observed bag location on trial t and the bucket position chosen on the previous trial:

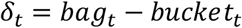

This approach allows us to characterize updating behavior without imposing assumptions from any specific computational model. In practice, rather than computing a ratio on each trial, we estimated learning rates at the block level by regressing belief updates onto prediction errors across trials within each block:

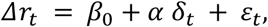

where *α* is the regression coefficient related to prediction errors (i.e., the learning rate for that block), *β*_0_ is the learning-independent block effect, and *ε*_*t*_ is noise.

### Psychological questionnaires

Following the behavioral task in Experiment 1, participants completed a battery of self-report questionnaires assessing psychiatric symptoms across multiple domains. We selected instruments to cover Internalizing symptoms (particularly anxiety and depression), impulsivity and behavioral dysregulation, and broader psychiatric characteristics in accordance with NIH reporting standards. The following instruments were administered: Generalized Anxiety Disorder Scale (GAD-7) ^47^, the Patient Health Questionnaire (PHQ-9) ^48^, the Penn State Worry Questionnaire (PSWQ-8) ^49^, the Intolerance of Uncertainty Scale (IUS-12) ^50^, and the Depression, Anxiety, and Stress Scale (DASS-21) ^51^, the Barratt Impulsiveness Scale–Brief (BIS-8) ^52^, the Gambler’s Belief Questionnaire (GBQ-21) ^53^, Compulsive Buying Scale (CBS-7) ^54^, Problem Video Game Playing Scale (PVP-9) ^55^, the Lie-Bet Screening Tool for Problem Gambling (LBT) ^56^, the Yale-Brown Obsessive-Compulsive Scale modified for compulsive buying (YBOC-CB-10) ^57^, the Community Assessment of Psychic Experiences Positive Scale (CAPE-15) ^58^, the Eating Attitudes Test (EAT-26) ^59^, the Alcohol Use Disorders Identification Test (AUDIT-10) ^60^, and DSM-5-TR Self-Rated Level 1 Cross-Cutting Symptom Measure—Adult (DSM-23) ^61^. In compliance with NIH requirements, participants also completed the World Health Organization Disability Assessment Schedule (WHODAS 2.0) ^62^.

In Experiment 2, we administered only the questionnaires that loaded most strongly onto the two latent factors identified in Experiment 1, allowing us to compute comparable factor scores with reduced participant burden. For Internalizing coverage, we preserved the GAD-7, PSWQ-8, and IUS-12. For Externalizing coverage, we retained the GBQ-21, CBS-7, PVP-9, LBT, YBOC-CB-10, and AUDIT-10. These measures were selected for their robust loadings onto the two latent psychiatric factors identified in the Experiment 1 factor analysis, ensuring continuity in the primary transdiagnostic dimensions across experiments.

### Factor analysis

We conducted exploratory factor analysis (EFA) on questionnaire responses to identify underlying latent factors capturing transdiagnostic psychiatric variation and to reduce collinearity among questionnaire scales. Factor analysis was performed using Maximum Likelihood Estimation (MLE), implemented via MATLAB’s factoran function. Given anticipated correlations between latent psychiatric traits, we used oblique (promax) rotation, which allows factors to correlate. In Experiment 1, responses from 15 questionnaires (188 items total) were analyzed. The optimal factor solution was determined by examining the scree plot per Cattell’s criterion ^63^, which showed a notable drop in eigenvalues after two factors. A two-factor structure was therefore retained, capturing distinct yet correlated psychiatric dimensions. For Experiment 2, we applied the same approach to a reduced set of questionnaires (9 questionnaires, 86 items). To improve factor stability, we pooled responses for each subject across both experimental sessions (172 items total per subject).

We conducted exploratory factor analysis (EFA) on questionnaire responses to reduce collinearity among questionnaire scales and identify underlying latent factors capturing transdiagnostic psychiatric variation. Factor analysis was performed using Maximum Likelihood Estimation (MLE) implemented via MATLAB’s factoran function. Considering anticipated correlations between latent psychiatric traits, we employed an oblique (promax) rotation to enhance interpretability. In Experiment 1, responses from 15 questionnaires (188 items total) were analyzed. The optimal factor solution was determined by examining the scree plot, identifying a notable decrease in eigenvalues after two factors. Thus, a two-factor structure was retained, capturing distinct yet correlated psychiatric dimensions. For Experiments 2, we applied the same analytical approach to a smaller set of questionnaires (9 questionnaires comprising 86 items). To gain better explanatory power for each factor, we used pooled survey responses for each subject across both sessions (172 items total for each subject).

### Factor labeling

As reported in the Results, the two retained factors were labeled based on the strongest and most consistent item-level loading patterns. Factor 1 (Internalizing) was primarily defined by high loadings from the PSWQ-8, with substantial contributions from the GAD-7, DASS-21, and PHQ-9. Factor 2 (Externalizing) was characterized by strong and consistent loadings from the GBQ-21, followed by meaningful contributions from the LBT and YBOC-CB-10. Factor labeling in Experiment 2 was determined in the same way. Factor 1 (Internalizing) was primarily defined by high loadings from the GAD-7, PSWQ-8, and IUS-12. Factor 2 (Externalizing) was primarily defined by loadings from the GBQ-21 and LBT, PVP-9, YBOC-CB-10, and CBS-7. Full factor loadings are reported in Supplementary Fig. 3.

### Learning rate effects and association with psychiatric factors

We used generalized linear models (GLMs) to examine whether individual differences in latent psychiatric dimensions were associated with distinct components of learning under uncertainty. Psychiatric dimensions were defined using factor scores derived from the exploratory factor analyses described above. For each participant, we extracted task-level learning rate components from the model-neutral analysis and used these as dependent variables in regression analyses with psychiatric factor scores as predictors.

We extracted three task-derived learning rate effects from the model-neutral analysis. The baseline learning rate (*B*) was defined as the average of block-level learning rates across the four blocks. The stochasticity effect (*S*) was defined as the difference in average learning rate between high- and low-stochasticity blocks. The volatility effect (*V*) was defined analogously, as the difference in average learning rate between high- and low-volatility blocks. Each effect was then used as the dependent variable in a separate GLM:

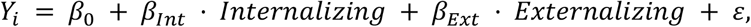

where *Y*_*i*_ denotes the learning rate effect of interest (*B, S*, or *V*), and Internalizing and Externalizing denote the two psychiatric factor scores. GLMs were fit using MATLAB’s glmfit function with an intercept term. This analysis tested whether psychiatric variation was associated with overall learning tendency (baseline) or with selective sensitivity to stochasticity or volatility, independently for each effect.

### CBF-HMM model

We implemented a Categorical Bayes Filter (CBF) ^31^ to analyze participants’ learning under binary outcomes without fixing volatility and stochasticity to known values in advance. The CBF uses the same binary state-space formulation as the standard hidden Markov model (HMM) but extends it by maintaining a joint posterior distribution over possible values of volatility and stochasticity, rather than treating these quantities as known. Unlike the particle filter implementation developed previously ^13^, which approximates this distribution via particle sampling with diffusion of uncertainty parameters across trials, the CBF approximates inference using a deterministic grid over candidate volatility and stochasticity values. This eliminates sampling-related randomness while preserving trial-by-trial inference about latent states and uncertainty.

We first describe the binary state-space model underlying the HMM, as described in our recent work ^13^. Let *x*_*t*_ ∈ {0,1} denote the hidden state on trial *t*, and let *o*_*t*_ ∈ {0,1} denote the corresponding binary outcome. The hidden state evolves according to a binary diffusion process governed by volatility *v*, such that

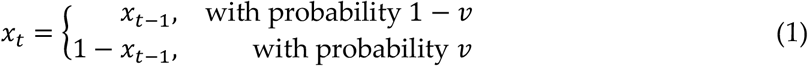

where 0 ≤ *v* ≤ 0.5. Thus, *v* determines the probability that the latent state switches from one trial to the next. When *v* = 0, the hidden state is perfectly stable, whereas *v* = 0.5 corresponds to maximal state uncertainty.

Outcomes are generated by corrupting the hidden state with stochasticity *s*, such that

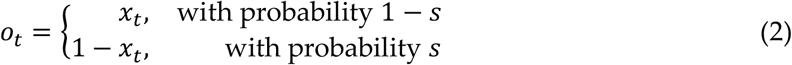

where 0 ≤ *s* ≤ 0.5. Larger values of *s* imply greater observational noise, such that the outcome becomes less reliable as an indicator of the hidden state.

Under this binary state-space model, the HMM provides optimal solution assuming known values of *v* and *s*. The posterior distribution over the hidden state is Bernoulli at every trial. Let *r*_*t*−1_ = *P* (*x*_*t*−1_ = 1 ∣ *o*_1:*t*−1_) denote the posterior probability that the latent state equals 1 on trial *t* − 1. For known values of *v* and *s*, inference proceeds using the standard HMM recursion. First, the belief before observing *o*_*t*_ is

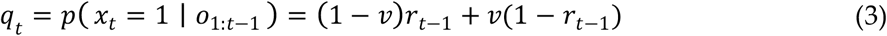

Second, the likelihood of the observed outcome under the hypothesis that *x*_*t*_ = 1 is

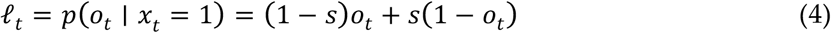

Applying Bayes’ rule then yields the posterior belief

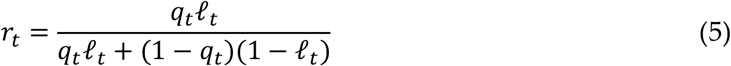

With initial condition *r*_0_ = 0.5, these equations provide exact HMM inference for known values of *v* and *s*.

To handle unknown *v* and *s*, we wrap the Categorical Bayes Filter (CBF) ^31^ around the HMM above, yielding a model we refer to as the CBF-HMM. Rather than fixing *v*and *s*in advance, the CBF-HMM maintains a categorical distribution over a deterministic grid of candidate parameter values. Within each block, the CBF represents uncertainty over volatility and stochasticity using two Beta distributions: one over candidate volatility values and one over candidate stochasticity values. Each Beta is parameterized by its mean *μ* ∈ (0, 1) and dispersion *η* ∈ (0, 0.5), with shape parameters *α* = *μ*/*η* and *β* = (1 − *μ*)/*η*. The four free parameters of the model are therefore *μ*_*v*_, *η*_*v*_, *μ*_*s*_, and *η*_*s*_: the means and dispersions of the two priors.

Each Beta distribution is discretized into *M* equally spaced quantiles *τ*_1_, … , *τ*_*M*_ ∈ [0.01, 0.99]. Candidate volatility and stochasticity values are obtained by inverse transform sampling from the corresponding Beta distributions:

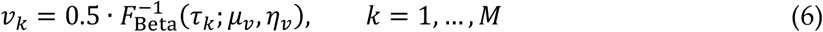

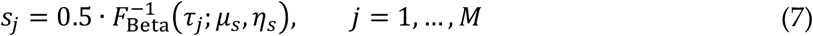

The factor of 0.5 rescales the Beta-distributed quantiles to the permissible parameter range 0,0.5. Taking the Cartesian product of the candidate volatility and stochasticity values yields a two-dimensional grid of *M*^2^ candidate *v, s* combinations. For brevity, we index these grid points by

*i* = 1, … , *M*^2^, with each *i* corresponding to a unique pair *k, j*; we write *v*^(*i*)^ and *s*^(*i*)^ for the volatility and stochasticity components of grid point *i*. The grid points are initialized with equal prior weight, 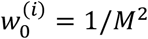, and the latent-state belief at every grid point is initialized as 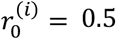 Thus, for each block the model is initialized with no prior preference for either hidden state or no prior bias toward any particular (*v, s*) combination.

For each grid point *i*, the model maintains both a belief state 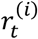 and an associated weight 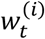. On each trial, each grid point generates its own predicted belief about the hidden state prior to observing the current outcome,

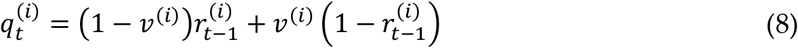

which implies a predictive probability for the outcome *o*_*t*_ being 1,

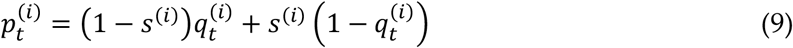

Given the observed outcome *o*_*t*_, the Bernoulli likelihood under grid point *i* is

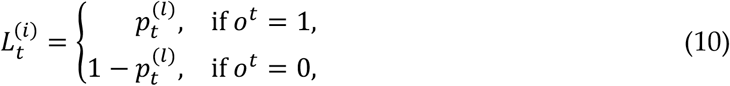

and the corresponding grid weight is updated by Bayesian importance weighting:

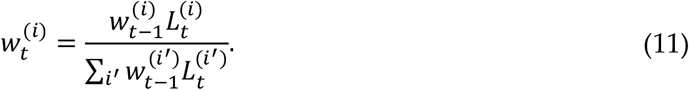

Candidate parameter combinations that better account for the observed outcome receive higher posterior weight over time.

After updating the weights, the hidden-state belief at each grid point is updated using the HMM equations with parameters (*v*^(*i*)^, *s*^(*i*)^). The model’s trial-wise latent prediction is then computed as the weighted average across all grid points,

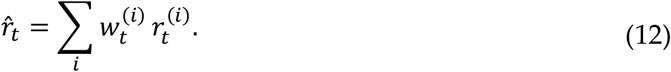

Observed choices were linked to latent beliefs through a separate response model. The CBF-HMM yields a trial-wise latent prediction 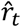, interpreted as the inferred probability that the hidden state equals 1 on trial *t*. However, choices may also reflect a tendency to repeat the previous response independently of latent-state inference and history of outcomes. To account for this, we augmented the model-derived prediction with a response-level stickiness parameter ρ. Specifically, choice probability was modeled in log-odds space as

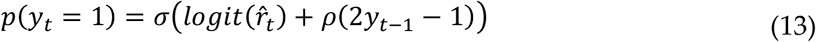

where *y*_*t*_ ∈ {0,1} denotes the observed choice on trial *t, σ*(*x*) = 1/(1 + *e*^−*x*^) is the logistic sigmoid function, and 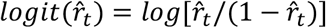. Under this formulation, positive values of *ρ* increase the tendency to repeat the previous choice, whereas negative values favor switching. The likelihood of the observed choice sequence was then computed as a Bernoulli likelihood under these response probabilities, separating latent-state inference from response biases. This formulation separates latent-state inference from response tendencies, allowing belief updating under uncertainty to be distinguished from motor or perseverative bias.

### Model fitting procedure

The CBF-HMM was fitted to each participant’s choice data via maximum a posteriori (MAP) estimation. Five free parameters were estimated per participant: the four CBF parameters governing the Beta priors over volatility and stochasticity (*μ*_*v*_, *η*_*v*_, *μ*_*s*_, *η*_*s*_), and the response stickiness parameter *ρ*. Optimization was performed in an unconstrained parameter space, with sigmoid transformations mapping each unconstrained parameter back to its admissible range in the generative model. Zero-mean Gaussian priors with variance 6.25 were placed on each unconstrained parameter, following the standard choice in the cbm toolbox ^64^ for parameters in the unit range. This pull estimates toward the center of the sigmoid, providing mild regularization without strongly constraining the fit.

MAP estimation was implemented via the Laplace approximation around the posterior mode, as provided by the cbm toolbox ^64^. For each participant, the negative log-posterior was minimized over the full sequence of choices across all blocks; trials with missing responses were excluded from likelihood evaluation. To mitigate sensitivity to local optima, optimization was run from 10 random initializations per participant, and the best-converged solution was retained.

To assess whether block-level learning rates can be reliably recovered from CBF-HMM fits, we performed a recovery analysis on *N* = 1000 synthetic datasets matching the trial structure of the binary experiments. For each simulated subject, true parameter values (*μ*_*v*_, *η*_*v*_, *μ*_*s*_, *η*_*s*_, *c*) were sampled from a Gaussian centered on the empirical mean and standard deviation of fitted parameters in the unconstrained parameter space. Belief trajectories were generated by running the CBF forward with these true parameters on the empirical outcome sequence, and binary choices were sampled trial-by-trial from a Bernoulli distribution with probabilities given by the response model. The same fitting procedure applied to human data was then applied to each simulated dataset. Recovery was quantified as the difference between true and recovered block-level learning rates across all simulated subjects (Supplementary Table 13, Supplementary Fig. 5).

### Learning rate effects and association with psychiatric factors (Experiment 2)

For Experiment 2, we extended the regression framework employed in Experiment 1 to capture learning components both shared across the two tasks and those varying with outcome valence. We defined average effects across the two tasks for baseline learning rate (*B*), stochasticity effect (*S*), volatility effect (*V*), and their interaction (*S* × *V*), together with corresponding valence contrast terms defined as the within-subject reward-minus-loss difference for each of these effects. For each psychiatric factor, we fit the combined regression model

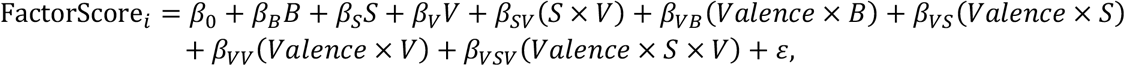

where *B, S, V*, and *S* × *V* denote the baseline, stochasticity, volatility, and interaction effects across the two tasks, and the Valence × terms denote the corresponding reward-minus-loss contrasts. Separate GLMs were fit for each factor.

For task-specific analyses, we additionally fit separate GLMs to each task individually without the Valence × terms. This allowed us to determine whether effects observed in the combined analysis were driven primarily by the reward task, the loss task, or both.

## Code and data availability

Analyses were conducted in MATLAB (R2023b) and Python (3.11.5). All experimental data and analysis code are publicly available online at https://github.com/piraylab/psyc-pathology-main.

## Acknowledgements

This work was supported by grants R21MH134217 from the National Institute of Mental Health. The authors declare no conflicts of interest.

## References

1. Browning, M., Behrens, T. E., Jocham, G., O’Reilly, J. X. & Bishop, S. J. Anxious individuals have difficulty learning the causal statistics of aversive environments. Nat. Neurosci. 18, 590–596 (2015).

2. Gagne, C., Dayan, P. & Bishop, S. J. When planning to survive goes wrong: Predicting the future and replaying the past in anxiety and PTSD. Curr. Opin. Behav. Sci. 24, 89–95 (2018).

3. Horga, G. & Abi-Dargham, A. An integrative framework for perceptual disturbances in psychosis. Nat. Rev. Neurosci. 20, 763–778 (2019).

4. Pulcu, E. & Browning, M. The Misestimation of Uncertainty in Affective Disorders. Trends Cogn. Sci. 23, 865–875 (2019).

5. Stephan, K. E., Baldeweg, T. & Friston, K. J. Synaptic plasticity and dysconnection in schizophrenia. Biol. Psychiatry 59, 929–939 (2006).

6. Zack, M., St George, R. & Clark, L. Dopaminergic signaling of uncertainty and the aetiology of gambling addiction. Prog. Neuropsychopharmacol. Biol. Psychiatry 99, 109853 (2020).

7. Dayan, P., Kakade, S. & Montague, P. R. Learning and selective attention. Nat. Neurosci. 3, 1218–1223 (2000).

8. Mathys, C., Daunizeau, J., Friston, K. J. & Stephan, K. E. A bayesian foundation for individual learning under uncertainty. Front. Hum. Neurosci. 5, 39 (2011).

9. Soltani, A. & Izquierdo, A. Adaptive learning under expected and unexpected uncertainty. Nat. Rev. Neurosci. 20, 635–644 (2019).

10. Piray, P. & Daw, N. D. A simple model for learning in volatile environments. PLoS Comput. Biol. 16, e1007963 (2020).

11. Piray, P. & Daw, N. D. A model for learning based on the joint estimation of stochasticity and volatility. Nat. Commun. 12, 6587 (2021).

12. Piray, P. & Daw, N. D. Computational processes of simultaneous learning of stochasticity and volatility in humans. Nat. Commun. 15, 9073 (2024).

13. Fang, X. & Piray, P. Inferring the causes of noise from binary outcomes: A normative theory of learning under uncertainty. 2026.03.01.708925 Preprint at 10.64898/2026.03.01.708925 (2026).

14. Deserno, L. et al. Volatility Estimates Increase Choice Switching and Relate to Prefrontal Activity in Schizophrenia. Biol. Psychiatry Cogn. Neurosci. Neuroimaging 5, 173–183 (2020).

15. Gagne, C., Zika, O., Dayan, P. & Bishop, S. J. Impaired adaptation of learning to contingency volatility in internalizing psychopathology. eLife 9, e61387 (2020).

16. Lawson, R. P., Mathys, C. & Rees, G. Adults with autism overestimate the volatility of the sensory environment. Nat. Neurosci. 20, 1293–1299 (2017).

17. Paliwal, S. et al. Subjective estimates of uncertainty during gambling and impulsivity after subthalamic deep brain stimulation for Parkinson’s disease. Sci. Rep. 9, 14795 (2019).

18. Piray, P., Ly, V., Roelofs, K., Cools, R. & Toni, I. Emotionally aversive cues suppress neural systems underlying optimal learning in socially anxious individuals. J. Neurosci. 1394–18 (2019) doi:10.1523/JNEUROSCI.1394-18.2018.

19. Powers, A. R., Mathys, C. & Corlett, P. R. Pavlovian Conditioning-Induced Hallucinations Result from Overweighting of Perceptual Priors. Science 357, 596–600 (2017).

20. Reed, E. J. et al. Paranoia as a deficit in non-social belief updating. eLife 9, (2020).

21. Ladouceur, R., Gosselin, P. & Dugas, M. J. Experimental manipulation of intolerance of uncertainty: A study of a theoretical model of worry. Behav. Res. Ther. 38, 933–941 (2000).

22. Luhmann, C. C., Ishida, K. & Hajcak, G. Intolerance of uncertainty and decisions about delayed, probabilistic rewards. Behav. Ther. 42, 378–386 (2011).

23. Beck, A. T. Depression: Causes and Treatment. (University of Pennsylvania Press, 1970).

24. Harlé, K. M., Guo, D., Zhang, S., Paulus, M. P. & Yu, A. J. Anhedonia and anxiety underlying depressive symptomatology have distinct effects on reward-based decision-making. PLOS ONE 12, e0186473 (2017).

25. Huang, H., Thompson, W. & Paulus, M. P. Computational Dysfunctions in Anxiety: Failure to Differentiate Signal From Noise. Biol. Psychiatry 82, 440–446 (2017).

26. Perandrés-Gómez, A., Navas, J. F., van Timmeren, T. & Perales, J. C. Decision-making (in)flexibility in gambling disorder. Addict. Behav. 112, 106534 (2021).

27. Vanes, L. D. et al. Contingency Learning in Alcohol Dependence and Pathological Gambling: Learning and Unlearning Reward Contingencies. Alcohol. Clin. Exp. Res. 38, 1602–1610 (2014).

28. Aylward, J. et al. Altered learning under uncertainty in unmedicated mood and anxiety disorders. Nat. Hum. Behav. 3, 1116–1123 (2019).

29. Frank, M. J., Seeberger, L. C. & O’reilly, R. C. By carrot or by stick: Cognitive reinforcement learning in parkinsonism. Science 306, 1940–1943 (2004).

30. Piray, P. The role of dorsal striatal D2-like receptors in reversal learning: A reinforcement learning viewpoint. J. Neurosci. 31, 14049–14050 (2011).

31. Chen, J. & Piray, P. Categorical Bayes Filtering for Computational Phenotyping in Adaptive Learning. 2026.05.14.725268 Preprint at 10.64898/2026.05.14.725268 (2026).

32. Satti, M. H. et al. Absence of Systematic Effects of Internalizing Psychopathology on Learning Under Uncertainty. eLife 14, (2025).

33. Izquierdo, A. & Jentsch, J. D. Reversal learning as a measure of impulsive and compulsive behavior in addictions. Psychopharmacology (Berl.) 219, 607–620 (2012).

34. Banerjee, N., Chen, Z., Clark, L. & Noël, X. Behavioural expressions of loss-chasing in gambling: A systematic scoping review. Neurosci. Biobehav. Rev. 153, 105377 (2023).

35. Robbins, T. W., Banca, P. & Belin, D. From compulsivity to compulsion: the neural basis of compulsive disorders. Nat. Rev. Neurosci. 25, 313–333 (2024).

36. Zhang, K. & Clark, L. Loss-chasing in gambling behaviour: neurocognitive and behavioural economic perspectives. Curr. Opin. Behav. Sci. 31, 1–7 (2020).

37. Zorowitz, S., Solis, J., Niv, Y. & Bennett, D. Inattentive responding can induce spurious associations between task behaviour and symptom measures. Nat. Hum. Behav. 7, 1667–1681 (2023).

38. Piray, P. Not all uncertainty is alike: volatility, stochasticity, and exploration. arXiv.org https://arxiv.org/abs/2605.19215v1 (2026).

39. Fan, H., Gershman, S. J. & Phelps, E. A. Trait somatic anxiety is associated with reduced directed exploration and underestimation of uncertainty. Nat. Hum. Behav. 7, 102–113 (2023).

40. Botvinick, M. & Toussaint, M. Planning as inference. Trends Cogn. Sci. 16, 485–488 (2012).

41. Bazarjani, A. & Piray, P. Default Feature Representations of the Cognitive Map. 2026.05.11.724436 Preprint at 10.64898/2026.05.11.724436 (2026).

42. Bazarjani, A. & Piray, P. Efficient Learning of Predictive Maps for Flexible Planning. 2026.02.11.705395 Preprint at 10.64898/2026.02.11.705395 (2026).

43. Piray, P. & Daw, N. D. Linear reinforcement learning in planning, grid fields, and cognitive control. Nat. Commun. 12, 4942 (2021).

44. Piray, P. & Daw, N. D. Reconciling flexibility and efficiency: medial entorhinal cortex represents a compositional cognitive map. Nat. Commun. 16, 7444 (2025).

45. Leeuw, J. R. de, Gilbert, R. A. & Luchterhandt, B. jsPsych: Enabling an Open-Source Collaborative Ecosystem of Behavioral Experiments. J. Open Source Softw. 8, 5351 (2023).

46. Zorowitz, S. & Bennett, D. nivlab/nivturk: Prolific v1.2. Zenodo 10.5281/zenodo.6609218 (2022).

47. Spitzer, R. L., Kroenke, K., Williams, J. B. W. & Löwe, B. A brief measure for assessing generalized anxiety disorder: The GAD-7. Arch. Intern. Med. 166, 1092–1097 (2006).

48. Kroenke, K., Spitzer, R. L. & Williams, J. B. W. The PHQ-9. J. Gen. Intern. Med. 16, 606–613 (2001).

49. Meyer, T. J., Miller, M. L., Metzger, R. L. & Borkovec, T. D. Development and validation of the Penn State Worry Questionnaire. Behav. Res. Ther. 28, 487–495 (1990).

50. Carleton, R. N., Norton, M. A. P. J. & Asmundson, G. J. G. Fearing the unknown: A short version of the Intolerance of Uncertainty Scale. J. Anxiety Disord. 21, 105–117 (2007).

51. Lovibond, S. H. & Lovibond, P. F. Depression Anxiety Stress Scales. 10.1037/t01004-000 (2011).

52. Steinberg, L., Sharp, C., Stanford, M. S. & Tharp, A. T. New tricks for an old measure: the development of the Barratt Impulsiveness Scale-Brief (BIS-Brief). Psychol. Assess. 25, 216–226 (2013).

53. Steenbergh, T. A., Meyers, A. W., May, R. K. & Whelan, J. P. Development and validation of the Gamblers’ Beliefs Questionnaire. Psychol. Addict. Behav. 16, 143–149 (2002).

54. Faber, R. J. & O’Guinn, T. C. A Clinical Screener for Compulsive Buying. J. Consum. Res. 19, 459–469 (1992).

55. Tejeiro Salguero, R. A. & Morán, R. M. B. Measuring problem video game playing in adolescents. Addict. Abingdon Engl. 97, 1601–1606 (2002).

56. Johnson, E. E. et al. The Lie/Bet Questionnaire for screening pathological gamblers. Psychol. Rep. 80, 83–88 (1997).

57. Monahan, P., Black, D. W. & Gabel, J. Reliability and validity of a scale to measure change in persons with compulsive buying. Psychiatry Res. 64, 59–67 (1996).

58. Capra, C., Kavanagh, D. J., Hides, L. & Scott, J. Brief screening for psychosis-like experiences. Schizophr. Res. 149, 104–107 (2013).

59. Garner, D. M., Olmsted, M. P., Bohr, Y. & Garfinkel, P. E. The eating attitudes test: Psychometric features and clinical correlates. Psychol. Med. 12, 871–878 (1982).

60. Saunders, J. B., Aasland, O. G., Amundsen, A. & Grant, M. Alcohol consumption and related problems among primary health care patients: WHO collaborative project on early detection of persons with harmful alcohol consumption--I. Addict. Abingdon Engl. 88, 349–362 (1993).

61. Association, A. P. Diagnostic and Statistical Manual of Mental Disorders, 5th Edition: DSM-5. (American Psychiatric Publishing, Washington, D.C, 2013).

62. Axelsson, E., Lindsäter, E., Ljótsson, B., Andersson, E. & Hedman-Lagerlöf, E. The 12-item Self-Report World Health Organization Disability Assessment Schedule (WHODAS) 2.0 Administered Via the Internet to Individuals With Anxiety and Stress Disorders: A Psychometric Investigation Based on Data From Two Clinical Trials. JMIR Ment. Health 4, e58 (2017).

63. Cattell, R. B. The Scree Test For The Number Of Factors. Multivar. Behav. Res. 1, 245–276 (1966).

64. Piray, P., Dezfouli, A., Heskes, T., Frank, M. J. & Daw, N. D. Hierarchical Bayesian inference for concurrent model fitting and comparison for group studies. PLOS Comput. Biol. 15, e1007043 (2019).

