## Supplementary Information for "Dissociating volatility and stochasticity reveals transdiagnostic computational signatures of psychopathology"

| Sex | Experiment 1 $N$ | Experiment 2 $N$ |
|---|---|---|
| Female | 1323 | 361 |
| Male | 1144 | 351 |
| Other | 51 | 9 |
| Unknown | 14 | 2 |

| Age | Experiment 1 $N$ | Experiment 2 $N$ |
|---|---|---|
| 19-35 | 1186 | 230 |
| 36-50 | 885 | 306 |
| 51-64 | 359 | 137 |
| 65+ | 101 | 50 |
| Unknown | 1 | 0 |

| Experiment 1 Ethnicity × Race | White | Black or African American | Asian | American Indian or Alaska Native | Native Hawaiian or Other Pacific Islander | Other | Unknown |
|---|---|---|---|---|---|---|---|
| Not Hispanic or Latino | 1489 | 266 | 214 | 10 | 5 | 13 | 8 |
| Hispanic or Latino | 329 | 28 | 7 | 13 | 2 | 65 | 18 |
| Unknown | 33 | 12 | 0 | 1 | 0 | 2 | 17 |

| Experiment 2 Ethnicity × Race | White | Black or African American | Asian | American Indian or Alaska Native | Native Hawaiian or Other Pacific Islander | Other | Unknown |
|---|---|---|---|---|---|---|---|
| Not Hispanic or Latino | 460 | 75 | 68 | 2 | 0 | 2 | 4 |
| Hispanic or Latino | 81 | 4 | 2 | 4 | 0 | 7 | 2 |
| Unknown | 4 | 2 | 0 | 0 | 0 | 2 | 4 |

**Supplementary Table 1. Demographic characteristics of the Experiment 1 ($N = 2532$) and Experiment 2 ($N = 723$) sample.**

| | $\delta$ | $\delta \times S$ | $\delta \times V$ | $\delta \times S \times V$ | $S$ | $V$ | $S \times V$ | $I$ |
|---|---|---|---|---|---|---|---|---|
| Coeff. | 1.344 | -0.057 | 0.029 | -0.003 | -0.182 | 0.654 | 0.204 | 1.185 |
| S.E.M. | 0.006 | 0.003 | 0.003 | 0.002 | 0.015 | 0.017 | 0.015 | 0.024 |
| t-value | 207.961 | -22.057 | 10.721 | -1.041 | -11.777 | 38.937 | 13.314 | 49.175 |
| p-value | <0.001 | <0.001 | <0.001 | 0.298 | <0.001 | <0.001 | <0.001 | <0.001 |

**Supplementary Table 2. Model-neutral regression analysis of the bird task data ($N = 2532$).** This table presents the regression results for all regressors in the model-neutral analysis. Columns list each regressor together with its coefficient estimate (Coeff.), standard error of the mean (S.E.M.), t-value, and p-value from two-sided t-tests across participants. The term $\delta$ is defined as the observation minus the choice, $\delta_t = bag_t - bucket_t$. The regression also yielded simple-effect terms for $S$, $V$, and $S \times V$, regardless of learning and prediction errors, which are not predicted by the joint-inference framework and are not interpreted further.

| Group | Stochasticity-blind | Intact | Volatility-blind |
|---|---|---|---|
| Mean | 38.163 | 38.894 | 38.911 |
| S.E.M. | 0.605 | 0.539 | 0.731 |
| One-way ANOVA | $F(2,1187) = 0.448, \quad P = 0.639$ | | |

**Supplementary Table 3. Age across the three computational phenotypes defined by critical-value grouping in Experiment 1.**

| Baseline LR | *Internalizing* | *Externalizing* | $I$ |
|---|---|---|---|
| Coeff. | 0.083 | -0.001 | 0 |
| S.E.M. | 0.023 | 0.023 | 0.020 |
| t-value | 3.655 | -0.045 | 0 |
| p-value | <0.001 | 0.964 | 1 |

| Volatility Effect | *Internalizing* | *Externalizing* | $I$ |
|---|---|---|---|
| Coeff. | -0.016 | -0.074 | 0 |
| S.E.M. | 0.023 | 0.023 | 0.02 |
| t-value | -0.695 | -3.294 | 0 |
| p-value | 0.487 | 0.001 | 1 |

| Stochasticity Effect | *Internalizing* | *Externalizing* | $I$ |
|---|---|---|---|
| Coeff. | 0.018 | -0.027 | 0 |
| S.E.M. | 0.023 | 0.023 | 0.020 |
| t-value | 0.783 | -1.186 | 0 |
| p-value | 0.434 | 0.236 | 1 |

| Factor Score Diff | $B$ | $S$ | $V$ | $S \times V$ | $I$ |
|---|---|---|---|---|---|
| Coeff. | 0.139 | 0.062 | 0.101 | 0.029 | 0 |
| S.E.M. | 0.034 | 0.035 | 0.035 | 0.035 | 0.034 |
| t-value | 4.034 | 1.772 | 2.908 | 0.832 | 0 |
| p-value | <0.001 | 0.076 | 0.004 | 0.406 | 1 |

**Supplementary Table 4. Associations between model-neutral learning components and factor scores in Experiment 1.** Generalized linear regressions relating task-derived computational components into the two psychiatric factor scores (Internalizing and Externalizing). For each predictor, regression coefficient, standard error of the mean (S.E.M.), t-value, and p-value are reported. All regressors and factor scores were z-scored before analysis.

| | $\delta$ | $\delta \times S$ | $\delta \times V$ | $\delta \times S \times V$ | $S$ | $V$ | $S \times V$ | $I$ |
|---|---|---|---|---|---|---|---|---|
| Coeff. | 1.082 | -0.065 | 0.075 | -0.011 | 0.003 | 0.006 | -0.007 | -0.010 |
| S.E.M. | 0.017 | 0.002 | 0.004 | 0.001 | 0.001 | 0 | 0.001 | 0 |
| t-value | 63.893 | -27.363 | 19.932 | -13.244 | 4.93 | 42.774 | -7.362 | -55.062 |
| p-value | <0.001 | <0.001 | <0.001 | <0.001 | <0.001 | <0.001 | <0.001 | <0.001 |

**Supplementary Table 5. Regression decomposition of fitted CBF-HMM learning rates in the reward-based task of Experiment 2.** This table presents the regression results for the fitted CBF-HMM learning rates estimated from the reward-based task. Columns list each regressor together with its coefficient estimate (Coeff.), standard error of the mean (S.E.M.), t-value, and p-value from two-sided t-tests across participants. The term $\delta$ is defined as the observation minus the model-based belief, $\delta_t = o_t - r_{t-1}$. $S$ and $V$ denote the true stochasticity and true volatility of the environment, respectively, and $I$ denotes the intercept.

| | $\delta$ | $\delta \times S$ | $\delta \times V$ | $\delta \times S \times V$ | $S$ | $V$ | $S \times V$ | $I$ |
|---|---|---|---|---|---|---|---|---|
| Coeff. | 1.216 | -0.061 | 0.088 | -0.01 | 0 | 0.006 | -0.004 | -0.009 |
| S.E.M. | 0.016 | 0.002 | 0.004 | 0.001 | 0.001 | 0 | 0.001 | 0 |
| t-value | 76.163 | -25.333 | 22.846 | -12.724 | -0.564 | 44.905 | -3.833 | -52.581 |
| p-value | <0.001 | <0.001 | <0.001 | <0.001 | 0.573 | <0.001 | <0.001 | <0.001 |

**Supplementary Table 6. Regression decomposition of fitted CBF-HMM learning rates in the loss-based task of Experiment 2.** This table presents the regression results for the fitted CBF-HMM learning rates estimated from the Turtle task. Columns list each regressor together with its coefficient estimate (Coeff.), standard error of the mean (S.E.M.), t-value, and p-value from two-sided t-tests across participants. The term $\delta$ is defined as the observation minus the model-based belief, $\delta_t = o_t - r_{t-1}$. $S$ and $V$ denote the true stochasticity and true volatility of the environment, respectively, and $I$ denotes the intercept.

| | $Valence \times B$ | $Valence \times S$ | $Valence \times V$ | $Valence \times S \times V$ |
|---|---|---|---|---|
| Coeff. | -0.134 | -0.004 | -0.013 | -0.001 |
| S.E.M. | 0.017 | 0.003 | 0.004 | 0.001 |
| t-value | -7.998 | -1.267 | -3.22 | -0.596 |
| p-value | <0.001 | 0.206 | 0.001 | 0.551 |

**Supplementary Table 7. Regression decomposition of valence differences in fitted CBF-HMM learning rates (**reward−loss; $N = 723$**).** Values report coefficient estimates (Coeff.), standard error of the mean (S.E.M.), t-values, and p-values from two-sided t-tests across participants for the aligned valence contrast in learning rates from fitted model. Positive coefficients indicate stronger effects in reward- than in loss-based task, whereas negative coefficients indicate stronger effects in loss- than in reward-based task.

| Internalizing Factor | $B$ | $S$ | $V$ | $S \times V$ | $Valence \times B$ | $Valence \times S$ | $Valence \times V$ | $Valence \times S \times V$ | $I$ |
|---|---|---|---|---|---|---|---|---|---|
| Coeff. | 0.081 | -0.057 | 0.006 | -0.049 | -0.084 | 0.084 | 0.024 | -0.043 | 0 |
| S.E.M. | 0.039 | 0.051 | 0.040 | 0.051 | 0.038 | 0.049 | 0.040 | 0.048 | 0.037 |
| t-value | 2.090 | -1.119 | 0.162 | -0.959 | -2.207 | 1.729 | 0.605 | -0.888 | 0 |
| p-value | 0.037 | 0.264 | 0.871 | 0.338 | 0.028 | 0.084 | 0.545 | 0.375 | 1 |

| Externalizing Factor | $B$ | $S$ | $V$ | $S \times V$ | $Valence \times B$ | $Valence \times S$ | $Valence \times V$ | $Valence \times S \times V$ | $I$ |
|---|---|---|---|---|---|---|---|---|---|
| Coeff. | -0.009 | 0.026 | -0.126 | -0.028 | 0.052 | -0.021 | -0.036 | 0.047 | 0 |
| S.E.M. | 0.039 | 0.051 | 0.040 | 0.051 | 0.038 | 0.049 | 0.040 | 0.048 | 0.037 |
| t-value | -0.238 | 0.506 | -3.156 | -0.561 | 1.363 | -0.442 | -0.900 | 0.968 | 0 |
| p-value | 0.812 | 0.613 | 0.002 | 0.575 | 0.173 | 0.659 | 0.368 | 0.333 | 1 |

| Factor Score Diff | $B$ | $S$ | $V$ | $S \times V$ | $Valence \times B$ | $Valence \times S$ | $Valence \times V$ | $Valence \times S \times V$ | $I$ |
|---|---|---|---|---|---|---|---|---|---|
| Coeff. | 0.091 | -0.083 | 0.132 | -0.020 | -0.136 | 0.106 | 0.060 | -0.090 | 0 |
| S.E.M. | 0.064 | 0.084 | 0.065 | 0.083 | 0.062 | 0.080 | 0.066 | 0.079 | 0.061 |
| t-value | 1.420 | -0.99 | 2.021 | -0.244 | -2.176 | 1.324 | 0.917 | -1.131 | 0 |
| p-value | 0.156 | 0.322 | 0.044 | 0.807 | 0.030 | 0.186 | 0.359 | 0.259 | 1 |

**Supplementary Table 8. Associations between task-averaged components and the valence contrasts across the reward- and loss-based binary tasks of Experiment 2.** Generalized linear regressions relating factor scores to shared learning effects and valence contrasts are shown for the Internalizing factor, Externalizing factor, and their score difference. Predictors include the shared baseline learning component ($B$), stochasticity effect ($S$), volatility effect ($V$), interaction effect ($S \times V$), and valence interaction terms capturing reward–loss differences in these components ($Valence \times B$, $Valence \times S$, $Valence \times V$, and $Valence \times S \times V$); $I$ denotes the intercept. Positive valence coefficients indicate stronger effects in the reward- than in the loss-based task. All regressors and factor scores were z-scored before analysis.

| Internalizing Factor | $B$ | $S$ | $V$ | $S \times V$ | $I$ |
|---|---|---|---|---|---|
| Coeff. | 0.013 | -0.021 | 0.027 | -0.074 | 0 |
| S.E.M. | 0.038 | 0.051 | 0.040 | 0.050 | 0.037 |
| t-value | 0.346 | -0.403 | 0.684 | -1.477 | 0 |
| p-value | 0.730 | 0.687 | 0.494 | 0.140 | 1 |
| **Externalizing Factor** | $B$ | $S$ | $V$ | $S \times V$ | $I$ |
| Coeff. | 0.018 | 0.008 | -0.136 | 0.021 | 0 |
| S.E.M. | 0.038 | 0.051 | 0.040 | 0.050 | 0.037 |
| t-value | 0.467 | 0.147 | -3.436 | 0.416 | 0 |
| p-value | 0.640 | 0.883 | 0.001 | 0.677 | 1 |
| **Factor Score Diff** | $B$ | $S$ | $V$ | $S \times V$ | $I$ |
| Coeff. | -0.004 | -0.028 | 0.164 | -0.094 | 0 |
| S.E.M. | 0.062 | 0.084 | 0.065 | 0.082 | 0.061 |
| t-value | -0.072 | -0.336 | 2.506 | -1.156 | 0 |
| p-value | 0.942 | 0.737 | 0.012 | 0.248 | 1 |

**Supplementary Table 9. Associations between CBF-HMM learning effects and factor scores in the reward-based task of Experiment 2.** Standardized linear regressions relating latent factor scores to fitted CBF-HMM learning components are shown for the Internalizing factor, Externalizing factor, and their score difference. Predictors include the baseline learning component ($B$), stochasticity effect on learning ($S$), volatility effect on learning ($V$), and their interaction effect ($S \times V$); $I$ denotes the intercept. All regressors and factor scores were z-scored before analysis.

| Internalizing Factor | $B$ | $S$ | $V$ | $S \times V$ | $I$ |
|---|---|---|---|---|---|
| Coeff. | 0.111 | 0.104 | -0.01 | -0.005 | 0 |
| S.E.M. | 0.039 | 0.048 | 0.040 | 0.049 | 0.037 |
| t-value | 2.846 | 2.146 | -0.262 | -0.109 | 0 |
| p-value | 0.005 | 0.032 | 0.793 | 0.913 | 1 |
| **Externalizing Factor** | $B$ | $S$ | $V$ | $S \times V$ | $I$ |
| Coeff. | -0.043 | -0.034 | -0.084 | -0.052 | 0 |
| S.E.M. | 0.039 | 0.048 | 0.040 | 0.049 | 0.037 |
| t-value | -1.114 | -0.699 | -2.112 | -1.058 | 0 |
| p-value | 0.266 | 0.485 | 0.035 | 0.290 | 1 |
| **Factor Score Diff** | $B$ | $S$ | $V$ | $S \times V$ | $I$ |
| Coeff. | 0.154 | 0.138 | 0.073 | 0.047 | 0 |
| S.E.M. | 0.064 | 0.079 | 0.065 | 0.081 | 0.061 |
| t-value | 2.411 | 1.733 | 1.128 | 0.579 | 0 |
| p-value | 0.016 | 0.084 | 0.260 | 0.563 | 1 |

**Supplementary Table 10. Associations between CBF-HMM learning effects and latent factor scores in the loss-based task of Experiment 2.** Standardized linear regressions relating latent factor scores to fitted CBF-HMM learning components are shown for the Internalizing factor, Externalizing factor, and their difference score. Predictors include the baseline learning component ($B$), stochasticity effect on learning ($S$), volatility effect on learning ($V$), and interaction effect ($S \times V$); $I$ denotes the intercept. All regressors and factor scores were z-scored before analysis.

| Internalizing Factor | $B$ | $S$ | $V$ | $S \times V$ | $Age$ | $I$ |
|---|---|---|---|---|---|---|
| Coeff. | 0.024 | -0.02 | 0.015 | -0.067 | -0.172 | 0 |
| S.E.M. | 0.038 | 0.052 | 0.040 | 0.050 | 0.038 | 0.038 |
| t-value | 0.624 | -0.377 | 0.374 | -1.332 | -4.564 | 0 |
| p-value | 0.533 | 0.707 | 0.709 | 0.183 | <0.001 | 1 |
| **Externalizing Factor** | $B$ | $S$ | $V$ | $S \times V$ | $Age$ | $I$ |
| Coeff. | 0.031 | 0.004 | -0.158 | 0.023 | -0.117 | 0 |
| S.E.M. | 0.038 | 0.052 | 0.040 | 0.050 | 0.038 | 0.037 |
| t-value | 0.812 | 0.073 | -3.909 | 0.457 | -3.115 | 0 |
| p-value | 0.417 | 0.941 | <0.001 | 0.648 | 0.002 | 1 |

**Supplementary Table 11. Associations between CBF-HMM learning effects with age and factor scores in the reward-based task of Experiment 2.** Standardized linear regressions relating latent factor scores to fitted CBF-HMM learning components are shown for the Internalizing factor, Externalizing factor, and their score difference. Predictors include the baseline learning component ($B$), stochasticity effect on learning ($S$), volatility effect on learning ($V$), their interaction effect ($S \times V$), and age; $I$ denotes the intercept. All regressors and factor scores were z-scored before analysis.

| Internalizing Factor | $B$ | $S$ | $V$ | $S \times V$ | $Age$ | $I$ |
|---|---|---|---|---|---|---|
| Coeff. | 0.113 | 0.107 | -0.047 | 0.021 | -0.17 | 0 |
| S.E.M. | 0.039 | 0.049 | 0.040 | 0.050 | 0.038 | 0.037 |
| t-value | 2.867 | 2.17 | -1.161 | 0.41 | -4.527 | 0 |
| p-value | 0.004 | 0.030 | 0.246 | 0.682 | <0.001 | 1 |
| **Externalizing Factor** | $B$ | $S$ | $V$ | $S \times V$ | $Age$ | $I$ |
| Coeff. | -0.033 | -0.043 | -0.095 | -0.046 | -0.118 | 0 |
| S.E.M. | 0.040 | 0.050 | 0.041 | 0.051 | 0.038 | 0.038 |
| t-value | -0.84 | -0.872 | -2.347 | -0.901 | -3.106 | 0 |
| p-value | 0.401 | 0.383 | 0.019 | 0.368 | 0.002 | 1 |

**Supplementary Table 12. Associations between CBF-HMM learning effects with age and factor scores in the loss-based task of Experiment 2.** Standardized linear regressions relating factor scores to fitted CBF-HMM learning components are shown for the Internalizing factor, Externalizing factor, and their score difference. Predictors include the baseline learning component ($B$), stochasticity effect on learning ($S$), volatility effect on learning ($V$), their interaction effect ($S \times V$), and age; $I$ denotes the intercept. All regressors and factor scores were z-scored before analysis.

| | Block 1 | Block 2 | Block 3 | Block 4 |
|---|---|---|---|---|
| $\rho$ | 0.896 | 0.868 | 0.898 | 0.883 |

**Supplementary Table 13. Block-wise recovery of learning rates for the CBF-HMM.** Reported values summarize the correspondence between simulated and recovered learning rates for each of the four task blocks. For each block, the table reports Spearman's rank correlation coefficient ($\rho$) between true and recovered learning rates across simulated datasets. Recovery was high across all blocks, indicating strong rank-order identifiability of block-wise learning rates.

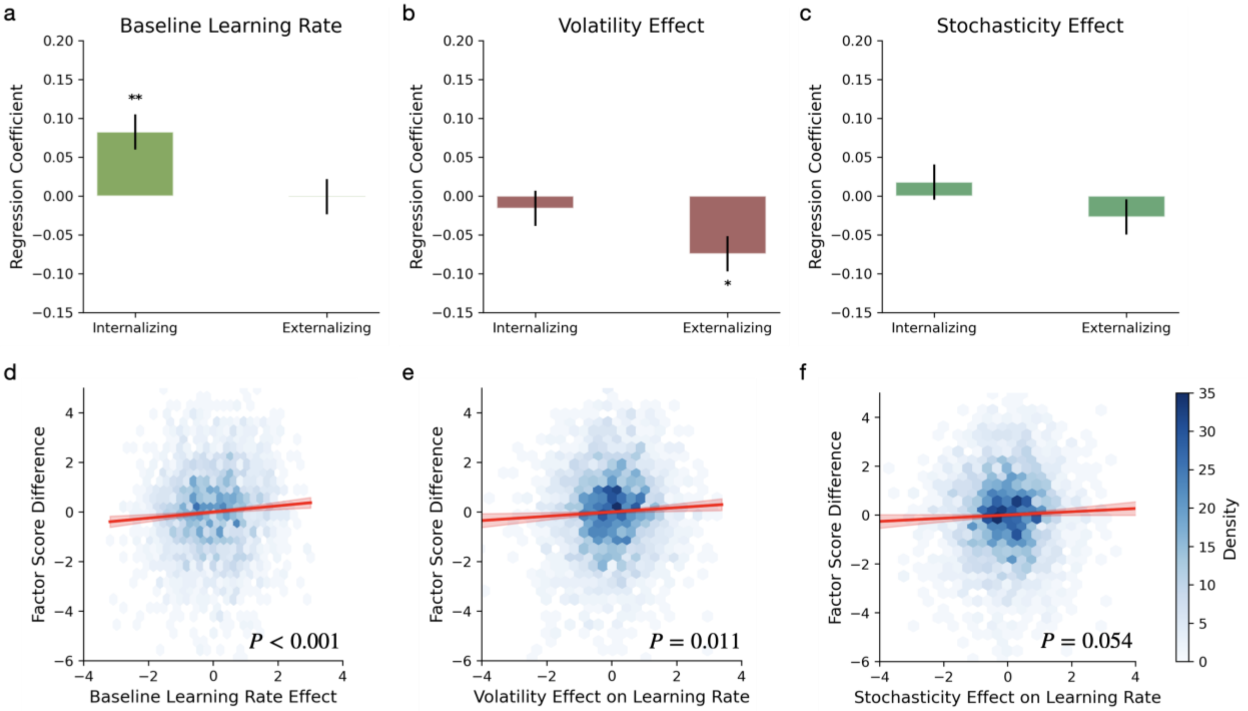


**Supplementary Fig. 1 Distinct dimensional associations between psychiatric factor scores and learning rate components.** a-c Regression coefficients from generalized linear models relating Experiment 1 factor scores to the baseline learning rate (a), volatility effect (b), and the stochasticity effect (c). Error bars indicate standard error of the mean. Asterisks denote statistical significance, with $P < 0.001$ (**) and $P < 0.05$ (*). d-f Hexagonal binned scatter plots showing the relationship between (Internalizing − Externalizing) score and bird task learning rate effects: baseline learning rate (d), volatility effect (e), and stochasticity effect (f). Red lines indicate linear regression fits; shaded regions indicate 95% confidence intervals. Color intensity represents participant density.

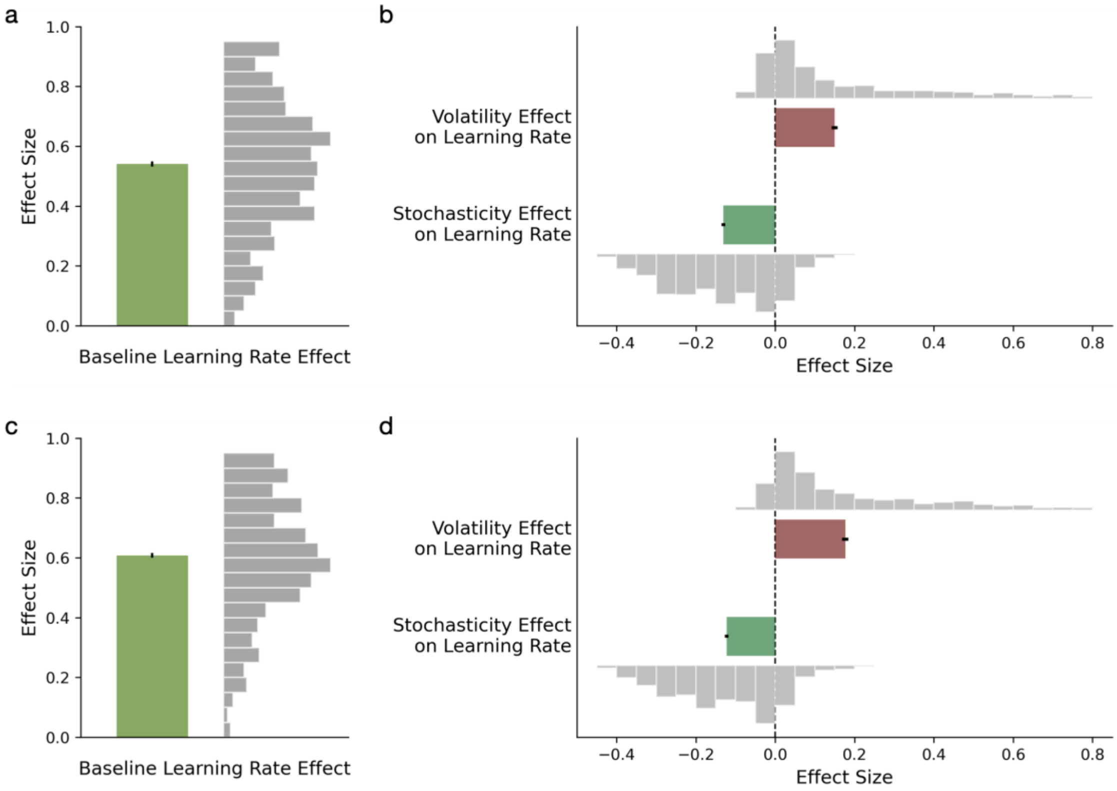


**Supplementary Fig. 2 CBF-HMM analysis of behavioral data in Experiment 2.** a–d, Mean baseline learning rate (a, c) and effects of volatility and stochasticity on learning rate (b, d) from CBF-HMM fits, for the reward-based (a, b) and loss-based (c, d) tasks; effects are computed as the difference between high- and low-condition blocks. Bars indicate the mean effect size, and error bars denote the standard error of the mean; adjacent histograms show the distribution of individual participant effect sizes.

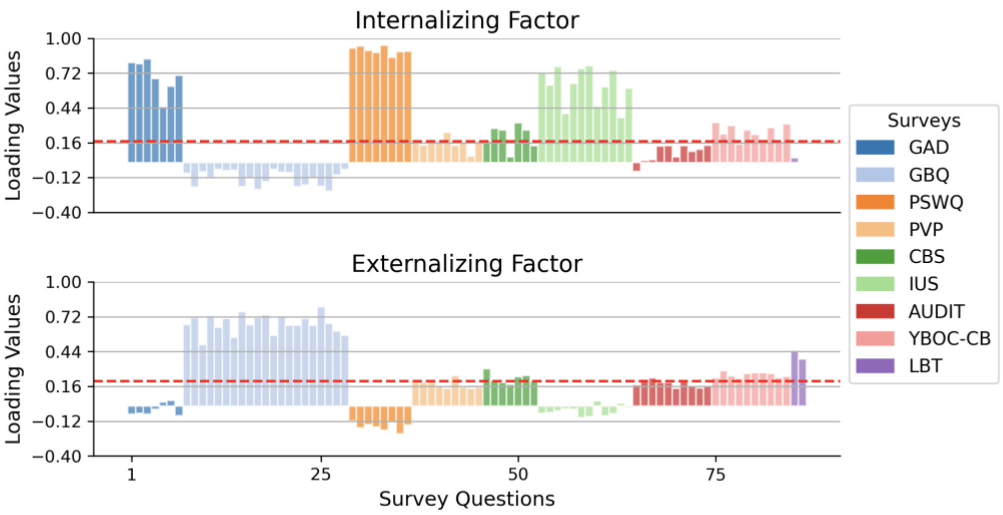


**Supplementary Fig. 3 Factor loadings in Experiment 2.** Factor loading values are shown for the two latent dimensions identified from exploratory factor analysis of questionnaire responses across the aligned reward- and loss-based sessions. Because the same participants completed the same questionnaire battery after both binary tasks, responses from matched participants were concatenated across sessions and a single shared factor model was fit to the combined item set, yielding one common latent structure for the binary-task sample. Questionnaire sources are differentiated by color across all items. The analysis revealed two primary factors that were readily interpretable as Internalizing and Externalizing dimensions.

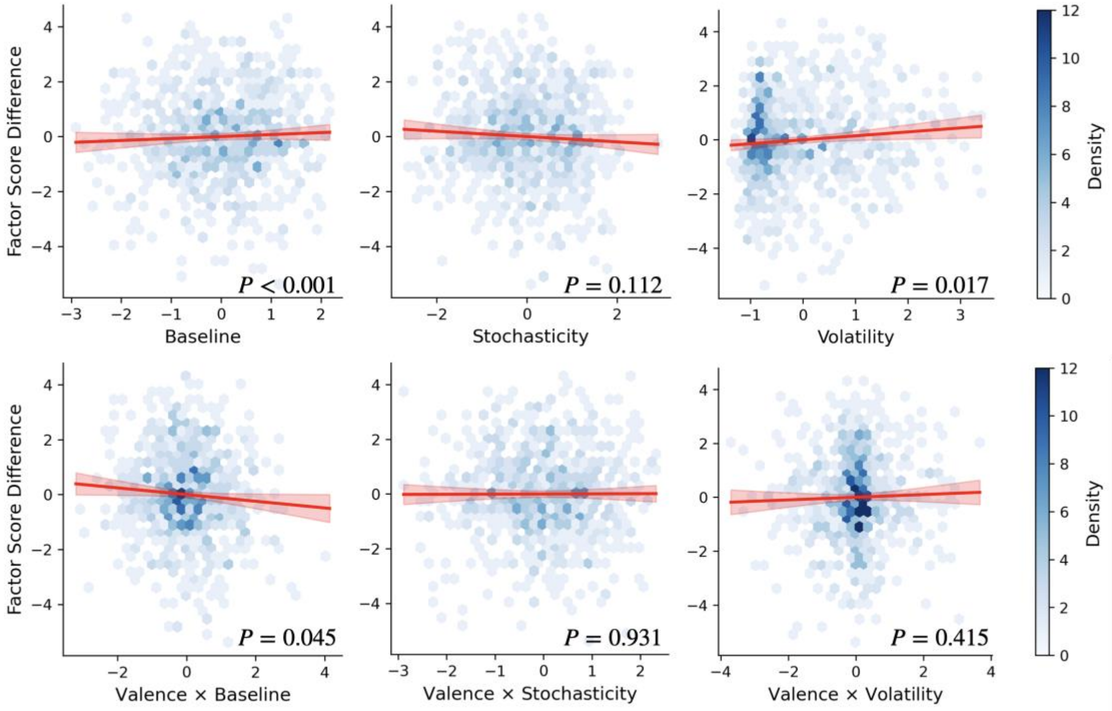


**Supplementary Fig. 4 Ordinary least squares associations of factor score difference between task-averaged components and the valence contrasts across the reward- and loss-based binary tasks of Experiment 2.** Hexagonal binned scatter plots show ordinary least squares (OLS) fits relating learning components from the reward- and loss-based tasks to psychiatric factor scores. Red lines indicate OLS regression fits, shaded regions indicate 95% confidence intervals, and color intensity reflects participant density.

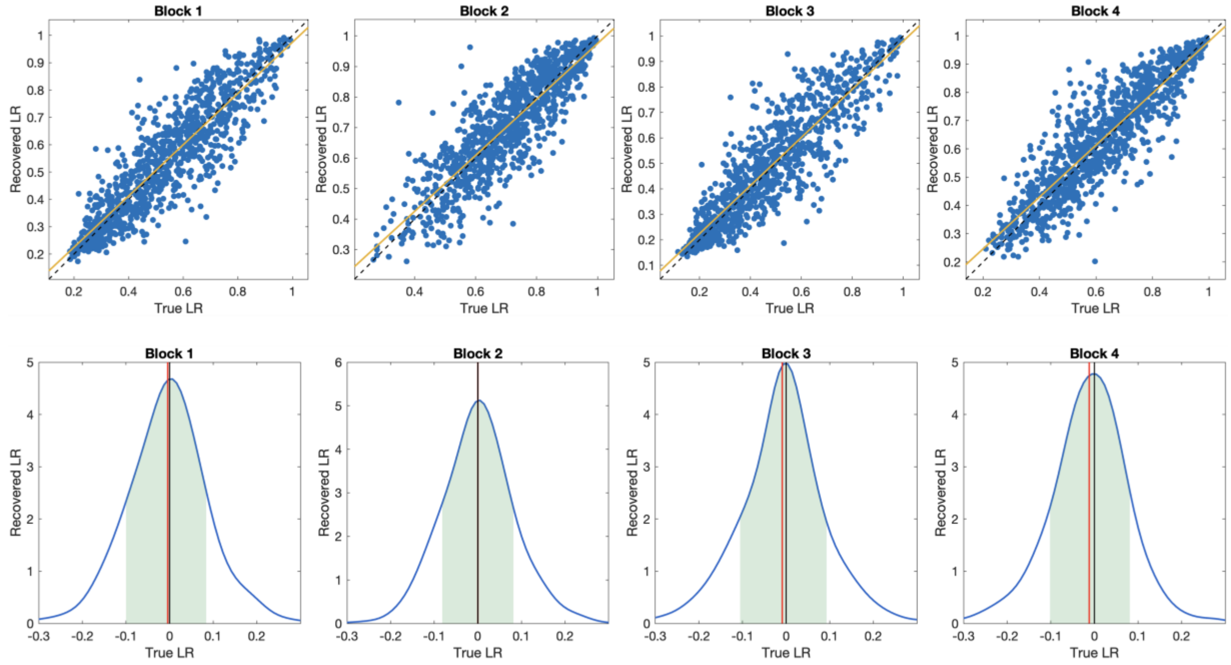


**Supplementary Fig. 5 Recovery of block-wise learning rates in the CBF-HMM recovery analysis.** Scatter plots show the relationship between simulated true and recovered learning rates for each of the four task blocks. The dashed diagonal indicates perfect recovery, and the solid fitted line shows the empirical linear relationship between true and recovered values. Across blocks, recovered learning rates tracked the simulated values closely, indicating good block-wise parameter identifiability. Kernel density plots of recovery errors for each block, defined as true minus recovered learning rate. Shaded regions denote ±1 standard deviation around the mean error, the red vertical line marks the mean error, and the black vertical line marks zero error. Error distributions were centered near zero across blocks, consistent with minimal systematic bias in learning-rate recovery.
